# Sulfur-mediated chalcogen versus hydrogen bonds in proteins: a seesaw effect in the conformational space

**DOI:** 10.1101/2022.03.14.484196

**Authors:** Vishal Annasaheb Adhav, Sanket Satish Shelke, P. Balanarayan, Kayarat Saikrishnan

**Author notes:** Correspondence and requests for materials should be addressed to K.S.

## Abstract

Divalent sulfur (S) form chalcogen bond (Ch-bond) *via* its σ–holes and hydrogen bond (H-bond) *via* its lone-pairs. Relevance of these interactions and their interplay for protein structure and function is unclear. Based on the analyses of the crystal structures of small organic/organometallic molecules and proteins, and their Molecular Electrostatic Surface Potential, we show that the reciprocity of the substituent-dependent strength of the σ–holes and lone-pairs correlate with the formation of either Ch-bond or H-bond. In proteins, disulfide-bonded cystine preferentially forms Ch-bond, metal-chelated cysteine forms H-bond, while methionine forms either of them with comparable frequencies. This has implications to the positioning of these residues and their role in protein structure and function. Computational analyses reveal that the S-mediated interactions stabilize protein secondary structures by mechanisms such as helix capping, protecting free β-sheet edges by negative-design, and augmenting the stability of β-turns. We find that Ch-bond can be as strong as H-bond. The study highlights the importance of S-mediated Ch-bond and H-bond for understanding protein folding and function, development of improved strategies for protein/peptide structure prediction and design, and structure-based drug discovery.

## Introduction

Non-covalent interactions are fundamental for protein folding, structural stability and function. Traditionally, hydrogen bonds (H-bonds), hydrophobic effects, electrostatic and van der Waals interactions are assumed to be the major drivers of protein folding and stability (Dill and MacCallum, 2012; Pace *et al.*, 2014). However, the essentiality of other weak interactions in sculpting protein structures is also being discovered. For example, the importance of weak H-bonds (such as C-H···O interaction), cation/anion-π and n→π^*^ interactions for the stability of protein structures is well understood (Derewenda *et al.*, 1995; Gallivan and Dougherty, 1999; Manikandan and Ramakumar, 2004; Bartlett *et al.*, 2010a; Lucas *et al.*, 2016; Newberry and Raines, 2019). Apart from C, O, N and H that form these non-covalent interactions, divalent sulfur (S) is present in methionines (Met-S^δ^) and cystines (Cys-S^γ^) of proteins. Sulfur has unique bonding properties that allows it to interact with both electrophiles and nucleophiles (Rosenfield *et al.*, 1977). Consequently, methionine and cystine are expected to contribute to distinct polar interactions in proteins, making it imperative to study them to better understand their structural properties and folding.

Though the polar bonding properties of S have long been known (Rosenfield *et al.*, 1977; Guru Row *et al.*, 1981), it has gained prominence in the field of organic molecules over the last decade (Andersen *et al.*, 2014; Beno *et al.*, 2015; Pascoe *et al.*, 2017; Motherwell *et al.*, 2018; Riwar *et al.*, 2018; Scilabra *et al.*, 2019; Haberhauer and Gleiter, 2020; Kolb *et al.*, 2020; Wang *et al.*, 2020). In contrast, these properties of S are often overlooked in the studies of proteins. For example, many standard textbooks of biochemistry categorize methionine as a non-polar hydrophobic amino acid (Lehninger, 2002; Berg, 2002). However, the lone-pairs of S can participate in H-bond formation with donor O or N (Zhou *et al.*, 2009; Rao *et al.*, 2015). It has also been shown recently that S of a disulfide bond can interact with backbone C=O through n→π^*^ interaction (Kilgore and Raines, 2018). Additionally, the divalent S has two electropositive regions along the extension of its two covalent bonds, referred to as σ–holes, which can interact with various nucleophiles (Murray *et al.*, 2012; Politzer *et al.*, 2014; Politzer *et al.*, 2017). The interaction made by a σ– hole of S with a nucleophile is categorized as a chalcogen bond (Ch-bond) (Aakeroy *et al.*, 2019).

Identified in many crystal structures of small, supra and biomolecules, chalcogen bonds have been shown to play important roles in self–assembly and catalysis of organic molecules (Pal and Chakrabarti, 2001; Iwaoka *et al.*, 2002; Garrett *et al.*, 2015; Benz *et al.*, 2017; Mahmudov *et al.*, 2017; Chen *et al.*, 2018; Lim and Beer, 2018; Vogel *et al.*, 2019). Ch-bonds have a specific directionality, and a recent spectroscopic study on thiophenes has shown that the bond can be as strong as a conventional H-bond (Pascoe *et al.*, 2017). However, unlike the latter, the strength of a Ch-bond is independent of solvent polarity (Pascoe *et al.*, 2017). The occurrence of Ch-bond in proteins has been documented previously (Pal and Chakrabarti, 2001; Iwaoka *et al.*, 2002; Iwaoka and Isozumi, 2012) and hypothesized to be functionally significant (Iwaoka *et al.*, 2002; Iwaoka and Isozumi, 2006). However, the precise role of Ch-bond in protein structure and its effect on protein stability and ligand binding have remained unaddressed.

Many functional groups in proteins contain both electrophilic and nucleophilic centers with which S can interact. Previous analysis of the crystal structures of organic molecules have shown that monovalent or divalent cations, H, C, etc., approach S roughly perpendicular to the X_1_-S-X_2_ plane (Rosenfield *et al.*, 1977), where X_1_ and X_2_ are substituents covalently bonded to S, presumably to interact with the lone-pairs (Figure 1a). On the other hand, anions, N, O, etc., approach S along the extension of the covalent bonds to interact with the σ–hole (Rosenfield *et al.*, 1977). Hence, the direction of approach of the functional groups with respect to S and the nature of the bond formed are interlinked, and could influence protein conformation.

**Figure 1.**
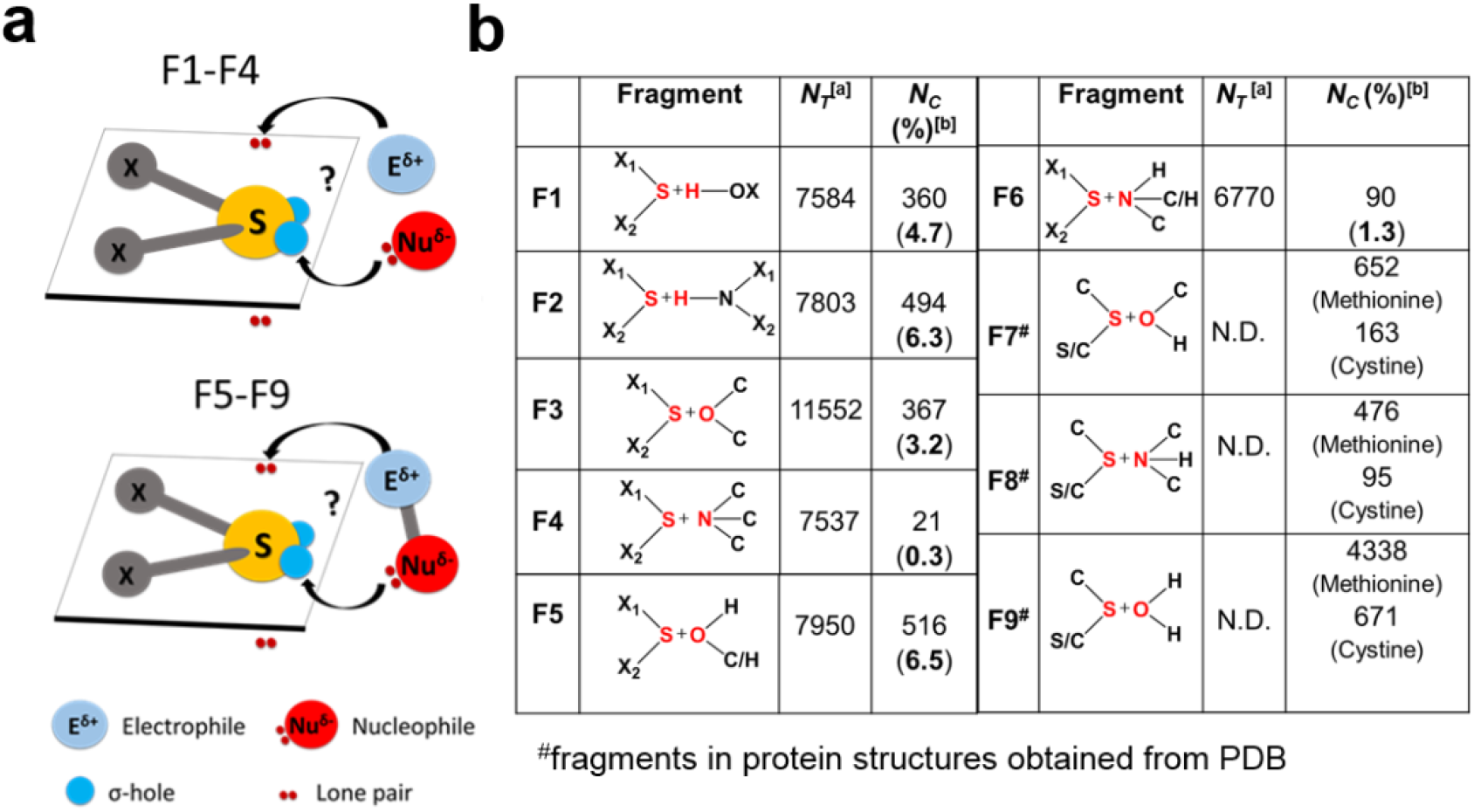
An approach of an electrophiles and nucleophiles towards divalent S. (**a**) Approach of an electrophile or a nucleophile (upper panel) and a covalently linked electrophile-nucleophile fragment (lower panel) towards S. (**b**) Fragments analyzed using the CSD and the PDB. *N_T_* is the total number of independent pair of fragments found in the CSD structures. *N_C_* is the total number of S···O or S···H-O and S···N or S···H-N contacts found in the CSD and the PDB, having distance between S and the atom in red less than the sum of their van der Waals radii. X_1_ and X_2_ are any element. Values in the parentheses are equal to *N_C_*/*N_T_* X 100, and represent the frequency of occurrence of the above-mentioned contacts in the CSD.

**Figure 2.**
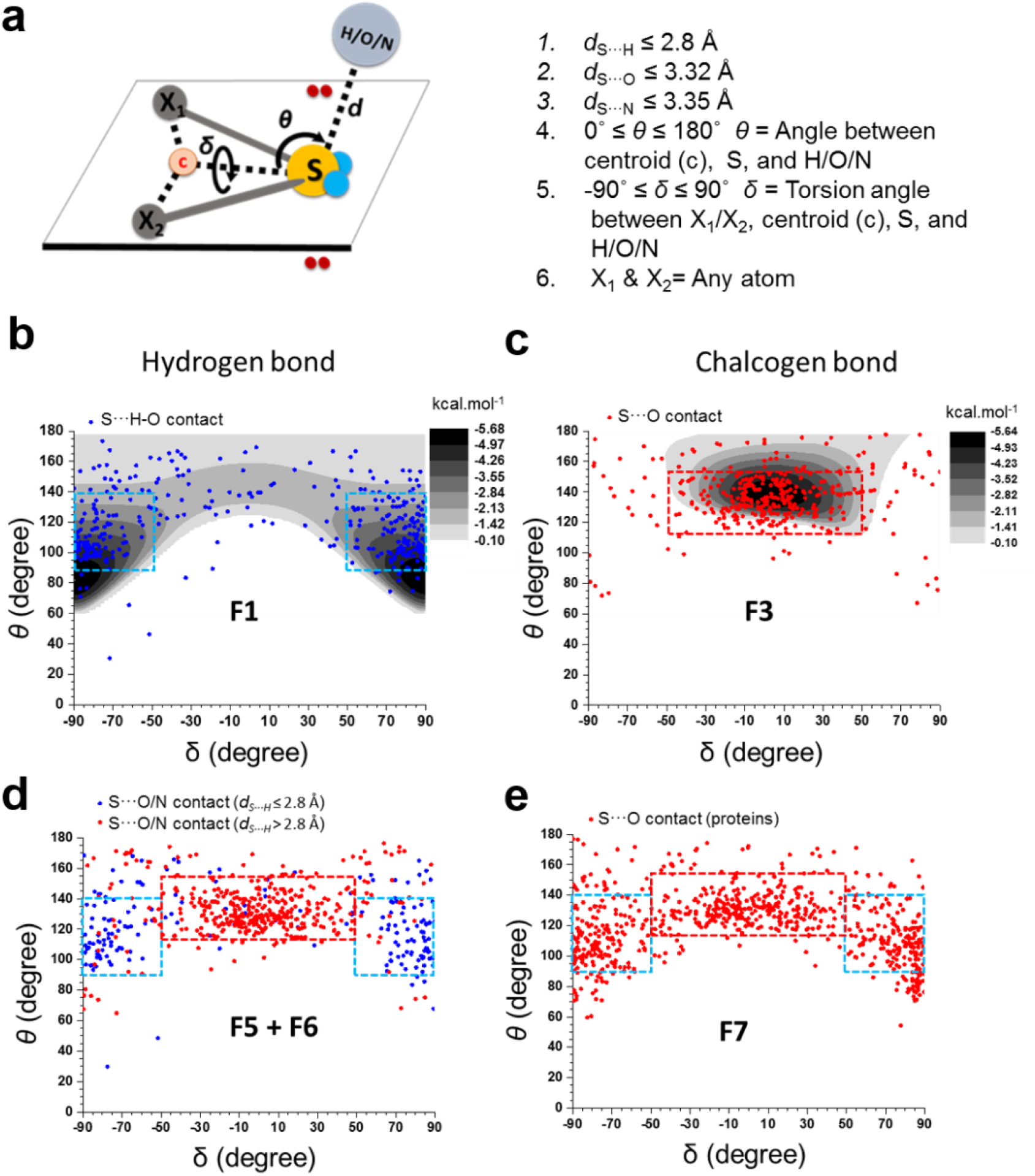
Nature of non-covalent interactions formed by S. **(a)** Definition of geometrical parameters *d*, *θ* and *δ*. **(b)** Mapping of *θ* and *δ* values of S···H-O contacts (blue dots) in F1 with computationally calculated ΔEs in the background (gray scale). (CH_3_)_2_S:OH_2_ complex was used as a model system to calculate ΔEs for F1; **(c)** S···O contacts (red dots) in F3. Cl(CH_3_)S:O(CH_3_)_2_ complex was used as the model system to calculate ΔEs in the background (gray scale). **(d)** S···O/N contacts with *d*_S···H_ ˃ 2.8 Å in red and those with *d*_S···H_≤ 2.8 Å in blue. **(e)** S···O contacts formed by methionine and cystine in F7.

This prompted us to also ask whether S forms a H-bond or a Ch-bond with a functional group having both electrophilic and nucleophilic centers (Figure 1a) and what determines the choice between them. Here, we have addressed the above questions through extensive computational, cheminformatics and bioinformatics analyses. The study, thus, unravels the underappreciated role of S-mediated interactions, in particular of Ch-bonds, in protein structure and stability.

## Methods

### Computational methods

All dimeric complexes *viz.* (CH_3_)_2_S:OH_2_, (CH_3_)_2_S:NH_3,_ Cl(CH_3_)S:O(CH_3_)_2_, and Cl(CH_3_)S:N(CH_3_)_3_ were optimized using the density functional M06 (Zhao and Truhlar, 2008) and the 6-31++G(2D,2P) basis set using the Gaussian09 suite of programs (Frisch *et al.*, 2009). In order to confirm that all these monomeric and dimeric optimized complexes are minima on the potential energy surface (PES), we performed Hessian evaluations with Int = ultrafine. It was confirmed that all the structures presented are minima on the PES from the positive eigenvalues of the Hessian. (CH_3_)_2_S:OH_2_ and (CH_3_)_2_S:NH_3_ complexes were used for PES scan to investigate energetically favorable region for S···H-O and S···H-N interactions, respectively. Similarly, PES scan were performed for Cl(CH_3_)S:O(CH_3_)_2_ for S···O and Cl(CH_3_)S:N(CH_3_)_3_ for S···N interactions. To carry out spherical energy scan of these complexes, *d* (distance between S and H/O/N) was kept constant, while *θ* (the imaginary angle between the centroid, c, of a triangle defined by C-S-X, S and H/O/N), and *δ* (the imaginary torsion angle between C, *c*, S and H/O/N) were varied (refer Supplementary Figure S1 for more details). Complexation energies (ΔEs) for different *θ* and *δ* values were calculated for all these complexes using the following equation (Řezáč and Hobza, 2016).

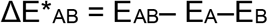

Here, E_A_ = ground state energy of monomer A, E_B_ = ground state energy of monomer B, E_AB_ = total energy of complex AB and ΔE*_AB_ = complexation energy of complex AB (*without correction for basis set superposition error). To calculate ΔEs, *δ* was varied from 90° to −90° (in steps of 2°) and *θ* from 60° to 178° (in steps of 2°). In the case of Cl(CH_3_)S:N(CH_3_)_3_, *θ* was varied from 90° to 178° when −40° ≤ *δ* ≤ 40° to avoid steric clashes among the atoms.

All the monomers mentioned in Figure 3a-b were optimized using M06 at 6-311++G(3DF,3PD). MESP topographical analyses characterizing the strength of lone-pairs (V_min_) of divalent S was carried out using the rapid topography mapping (Yeole *et al.*, 2012) as implemented in DAMQT (Kumar *et al.*, 2015). Texturing of MESP of molecules at defined density surface (V_s,max_) were carried out using Gaussian09.

**Figure 3.**
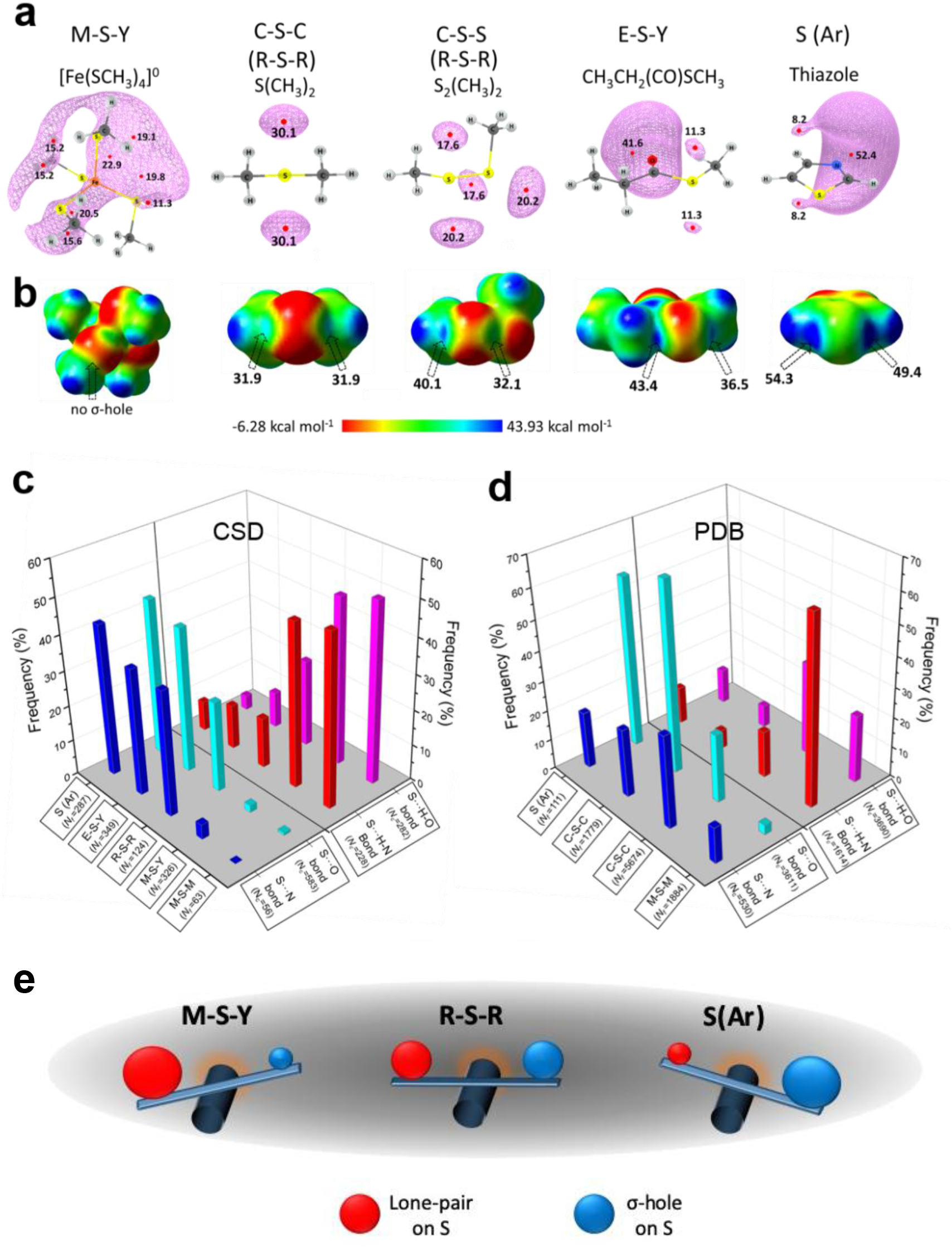
Rules for the formation of H- and Ch-bonds. **(a)** MESP minimum values, V_min_ in kcal.mol^−1^, represent the lone-pair regions of the S-containing monomers used in this study. **(b)** MESP map of the monomers with the two σ–holes marked by arrows and their magnitude, V_s,max_ in kcal.mol^−1^. **(c)** Histogram showing frequency formation of Ch-bond and H-bond in the CSD and **(d)** the PDB with different electronic environment of S. M = any metal; Y = any element except M; R = saturated C, H and S; E = any electron withdrawing group; and S(Ar)= Aromatic S. **(e)** Substituent-dependent seesaw change in the strength of the lone-pairs and σ–holes on S.

**Figure 4.**
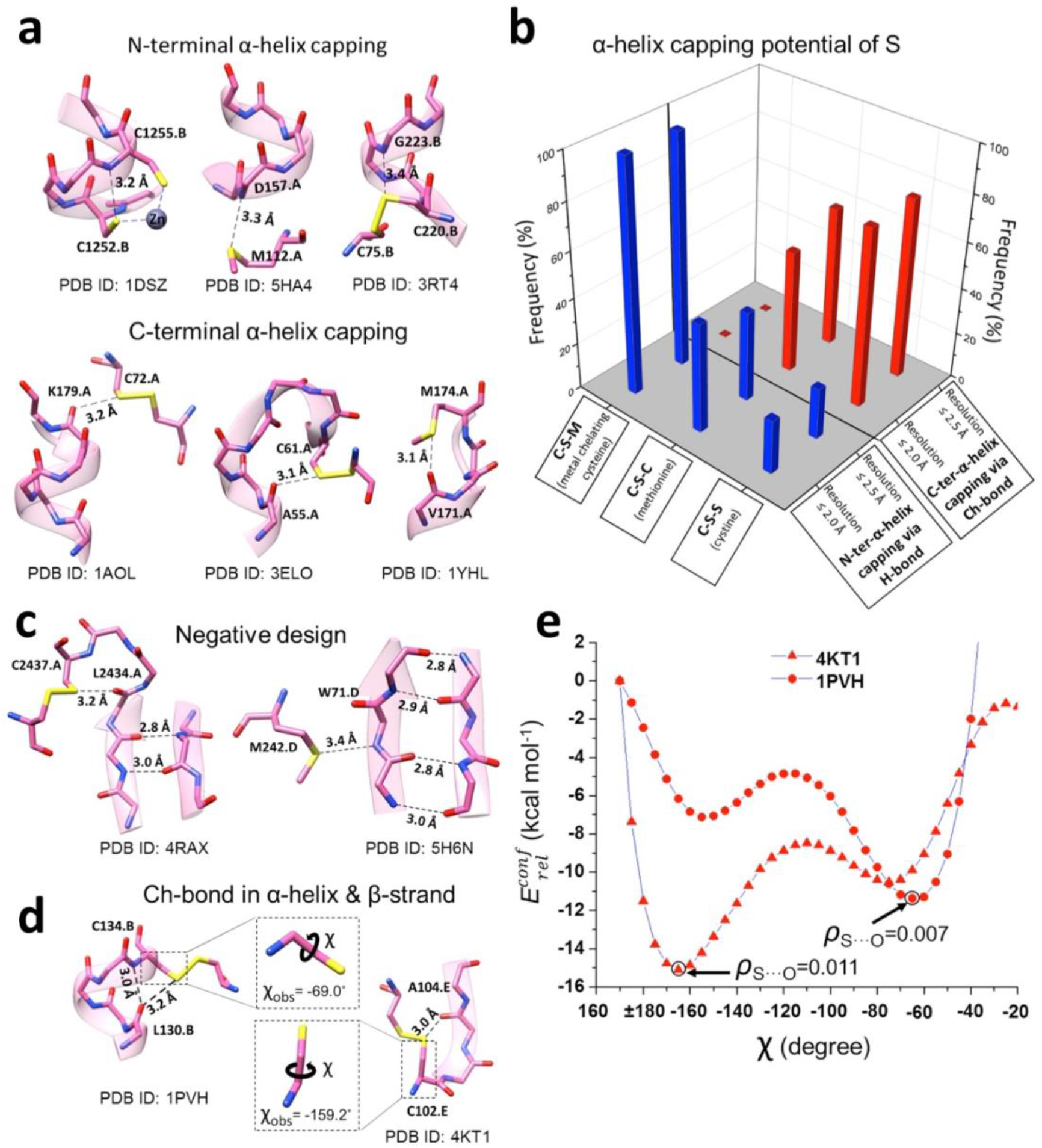
Stabilization of α-helices and β-sheets by S. (**a**) Representative examples of H-bond capping the N-terminus of α-helices, and of Ch-bond capping the C-terminus of α-helices. (**b**) Histogram showing the frequency of H-bond and Ch-bond interactions capping the N- and C-termini of α-helices by metal-chelating cysteine, methionine and cystine. (**c**) Representative examples of negative-design involving Ch-bond and H-bond. (**d**) Representative examples of Ch-bond found in α-helical and β-sheet regions. (**e**) The plot for conformational energies as a function of χ (E at χ = 170° was assigned as 0.0 kcal.mol^−1^). Values of ρ at BCP for S···O interaction for the most energetically favorable structures are given in au.

Model systems used to investigate the role of S···O interactions in α-helix is from the coordinates of PDB ID 1PVH, and 4KT1 for β-strand. All the side chain atoms in these fragments were deleted and C^β^ atom was replaced by H. The hydrogen atoms were added manually using Gaussview 5.0 program (Dennington *et al.*, 2009) following which partial energy minimization was carried out (all atoms other than hydrogen were frozen). The torsion angles (*χ*) between N, C^α^, C^β^ and S (Figure 5b) were varied for PES scan. A similar strategy was used to optimize the representative structures used for studying Cases 1 to 3 of CXXXXC motif (PDB IDs 2FD6 for Case 1, 3CEL for Case 2 and 1HTR for Case 3). AIM analysis was carried out using AIM2000 (König *et al.*, 2001) for the three structures.

**Figure 5.**
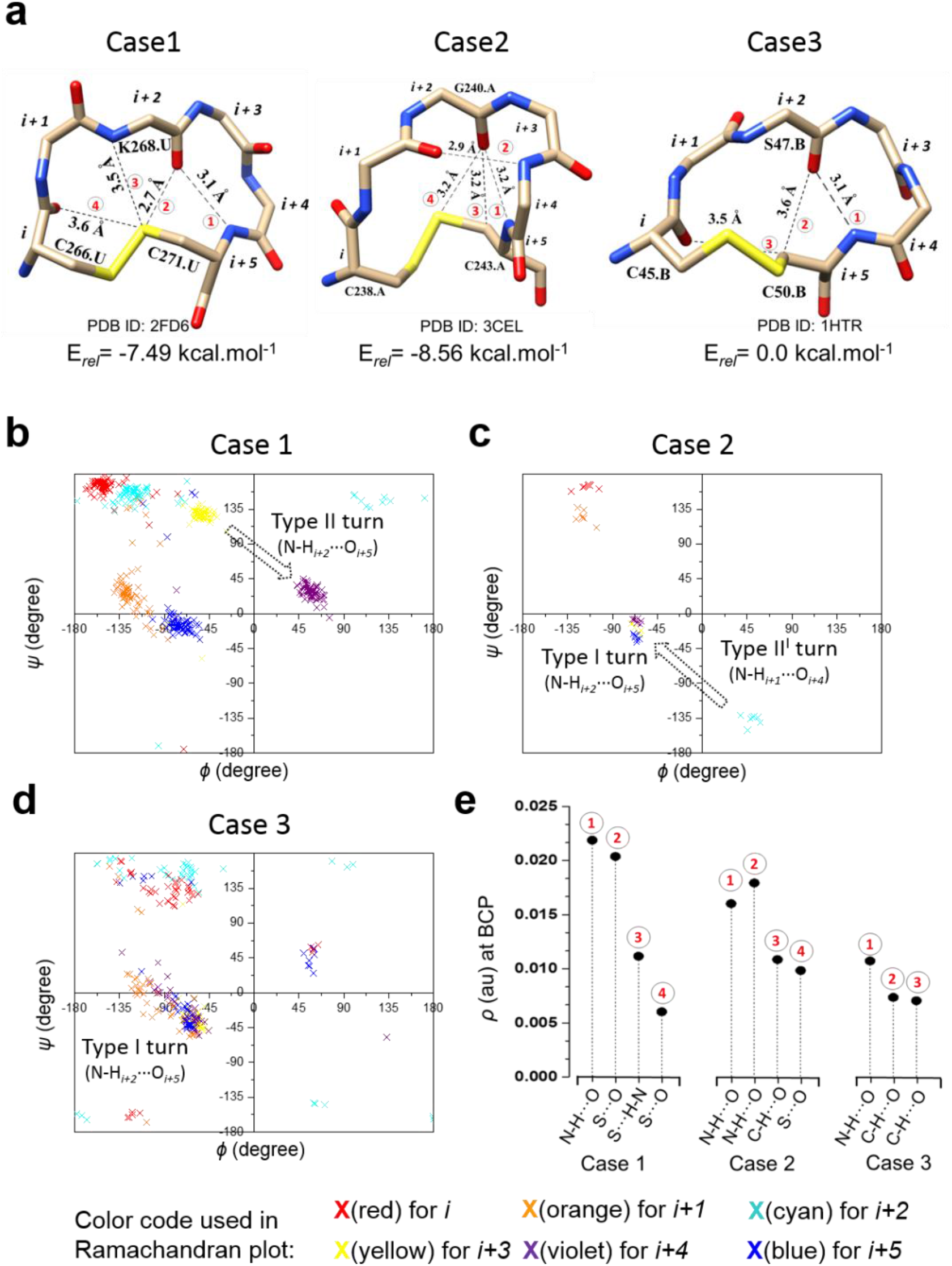
Contribution of Ch-bond to the stability of β-turns. (**a**) Representative examples of disulfide linked CXXXXC motif in protein structures. Left panel is Case 1 having a type II β-turn. Central panel is Case 2 having a type I and a type II’ β-turn. Right panel is Case 3 having a type I β-turn. Co-existing S···O, S···H-N, C···H-O and O···H-N interactions if present are marked by dotted lines and numbered in red in each panel. E_*rel*_ calculated for the three structures are given below the respective panel. (**b**-**d**) *Φ* and *Ψ* values of the residues from *i* to *i* + 5 in CXXXXC motifs belonging to Cases 1-3 shown in the Ramachandran plot. Structures satisfied the criteria of resolution *≤* 2.5 Å, *R_free_ ≤* 30% and pair-wise sequence identity ≤ 90% amongst them. (**e**) A plot showing *ρ* (kcal.mol^−1^) at BCP for interactions obtained using AIM for Cases 1-3.

### CSD analysis

Fragments provided in Figure 1b were queried in structural data retrieved from Cambridge Structural Database (CSD) (Groom and Allen, 2014) (version 5.39, Feb 2018) using ConQuest 1.21 (Bruno *et al.*, 2002). Our choice of fragments throughout this study was motivated by their relevance to proteins and their ligands. The following criteria available in ConQuest were used for the search: 1) 3D co-ordinates determined for all the atoms; 2) structures with crystallographic R factor ≤ 10%; 3) no disorder in crystallographic data; 4) no error in 3D atomic co-ordinates; and 5) no polymeric structures. Obtained data were further processed and analyzed using Mercury 3.10.1 (Bruno *et al.*, 2002). Searches were made for only intermolecular S···O/N and S···H-O/N contacts, where *d_S···O_* ≤ 3.32 Å (Bondi, 1964), *d_S···N_* ≤ 3.35 Å (Bondi, 1964) and *d_S···H_* ≤ 2.8 Å (Zhou *et al.*, 2009) (Supplementary Figure S2). The approach of H or O/N towards S in space was investigated using the 3D parameters facility provided in ConQuest 1.21. To segregate H- and Ch-bond based on their *θ* and *δ* values, we calculated the mean values for clusters in F1 and F2. The range of *θ* and *δ* defining H- and Ch-bond about the respective mean values were obtained by taking their mean ± standard deviation at 1 sigma of the calculated values for F1 and F2, respectively. The values for the limits were rounded off to the closest value that was a multiple of 5 (Supplementary Figure S3 for *θ* and S4 for *δ*). Mean of the angular values of *θ* was calculated from their modulus. This angular range of *θ* and *δ* for H- and Ch-bonds thus obtained were (see Results and Discussions for values) used throughout the study. All the plots were generated using OriginPro 9.0 (Seifert, 2014).

### PDB analyses

Protein structures determined using X-ray crystallography in the Protein Data Bank (PDB) (Rose *et al.*, 2017) were downloaded using PISCES (Wang and Dunbrack, 2005) on January 2018. Two sets of protein structures were generated using the following criteria. Set 1 contained protein structures having pairwise sequence identity ≤ 90%, *resolution* ≤ 2.0 and *R_free_* ≤ 25%. Set 2 contained protein structures having pairwise sequence identity ≤ 90%, *resolution* ≤ 2.5 Å and *R_free_* ≤ 30%. Coordinate files greater than 1 MB in size were excluded from both the sets. This resulted in Set 1 having 16851 structures and Set 2 having 25423 structures. These structures were analyzed using in-house scripts written in Python 3.7.1. Search for H-bonds and Ch-bonds were made using the criterion of *d* (Å) ≤ less than equal to sum of van der Waals radii of S and O/N (Bondi, 1964). To minimize the effect of structural constraints on the direction parameters, we excluded the contacts where the S and O/N were separated by less than 7 covalent bonds. The criteria for direction obtained using the CSD analysis was used to distinguish between H- and Ch-bond. Figures of protein structures were made using Chimera 1.13.1 (Pettersen *et al.*, 2004).

Search for metal-chelating cysteine used distance between S and metal to be within 1.9 to 2.8 Å. The distance range ensured that metals with different ionic radii were identified. The criteria for H-bond used was *d* ≤ 3.6 Å (Zhou *et al.*, 2009), 90°≤ *θ* ≤140°, −90°≤ *δ* ≤-50° or 50°≤ *δ* ≤90°. To understand the role of H-bond and Ch-bond in capping of α-helix, we searched for S···H-N (*d_S···N_* ≤ 3.6 Å, 90°≤ *θ* ≤140°, −90°≤ *δ* ≤−50° or 50°≤ *δ* ≤90°, and 120° ≤ *ζ*≤ 240°) and S···O (*d_S···O_* ≤ 3.32Å, 115°≤ *θ* ≤155° and −50°≤ *δ* ≤50°) interactions made by the peptide backbone with S. An additional criterion of an imaginary torsion angle *ζ* was applied to exclude structures where N-H was not pointing towards the lone-pair regions of S (refer Supplementary Figure S5 for more details). α-helices were identified using the header information in the PDB coordinate files. For N-terminal capping, we searched for S···H-N interactions where the amino group was of N_1_, N_2_ or N_3_ residue at helix terminus, as defined previously by Aurora and Rose (Aurora and Rose, 1998). While, for C-terminal capping, we searched for S···O interactions where the carbonyl O was of C_1_, C_2_ or C_3_ residue at helix terminus, as defined previously (Aurora and Rose, 1998).

To investigate S···H-N (*d_S···N_* ≤ 3.6 Å, 90°≤ *θ* ≤140°, −90°≤ *δ* ≤-50° or 50°≤ *δ* ≤90°and 120° ≤ *ζ*≤ 240°), S···O (*d_S···O_* ≤ 3.32Å, 115°≤ *θ* ≤155° and −50°≤ *δ* ≤50°) and S···N (*d_S···N_* ≤ 3.35Å, 115°≤ *θ* ≤155° and −50°≤ *δ* ≤50°) interactions in α-helices and β-strands, we obtained the information of secondary structures from the header in the PDB coordinate files. Only internal-helical residues were considered and the last four residues at the N- and the C-termini of the helix were excluded. In case of β-strands, all the residues belonging to the strand were considered.

To understand the role of S···O interaction in the CXXXXC motif, we searched disulfide linked cysteines separated by 4 intervening residues. The cyclic peptide was classified into four different groups namely, Case 1: Co-existing S···O bond between the S of *i* or *i*+5 cysteine and the carbonyl O of *i*+2 residue (*d_S···O_* ≤ 3.32Å, 115°≤ *θ* ≤155° and −50°≤ *δ* ≤50°), and H-bond between *i*+2 and *i*+5 residues (*d_O···N_* ≤ 3.5Å); Case 2: Co-existing S···O bond between the S of the *i* or *i*+5 residue and the carbonyl O of the *i*+2 residue, and H-bonds between *i*+2 and *i*+5 residues and *i*+1 and *i*+4 residues; Case 3: H-bond between *i*+2 and *i*+5 residues and absence of S···O bond between *i* or *i*+5 residue and *i*+2 residue; and Case 4: absence of any of the above mentioned non-covalent bonds.

## Results and Discussion

### Geometrical features that distinguish S-mediated Ch-bond from H-bond

As most structures in the PDB solved using X-ray crystallography do not have information of the position of hydrogen atoms, it is not trivial to identify if the non-covalent bond between S and O/N is H-bond or Ch-bond. An electrophile is expected to approach S along a direction different from a nucleophile, because the former interacts with the lone-pair while the latter interacts with the σ-hole (Figure 1a). To overcome the ambiguity due to unavailability of hydrogen atom coordinates, we devised a methodology based on the direction of approach of O or N towards S to distinguish between the H- and Ch-bonds. We first analyzed high-resolution crystal structures of organic molecules having experimentally determined hydrogen positions in the CSD to identify the preferred direction of approach of electrophiles towards S (**F1** and **F2**) to form H-bond or of nucleophiles (**F3** and **F4**) to form Ch-bond (Figure 1b).

*θ* and *δ* were chosen to obtain the preferred angular distribution for H- and Ch-bonds formed by S (Figure 2a). Potential Ch-bond between S and O/N required *d_S···O_* ≤ 3.32 Å or *d_S···N_* ≤ 3.35 Å (Bondi, 1964), while H-bond required *d_S···H_* ≤ 2.8 Å (Zhou *et al.*, 2009) (Figure 1c). The parameters measured for all such contacts in the CSD were plotted to illustrate the preferred range of distances and angles of these interactions (Figure 2b, e; Supplementary Table S1 and S2-S5). Use of these geometric parameters also facilitated a direct comparison of the angular distribution with the complexation energy ΔE (see Methods for definition) obtained from Potential Energy Surface (PES) scans at different values of *θ* and *δ* for the model systems (Supplementary Figure S1). ΔE was mapped on to the plot (Figure 2b,c; Supplementary Figure S6a).

Two distinct clusters were observed in the *θ*-*δ* plot for S···H-O contacts from (i) 90°≤ *θ* ≤140° and −50°≤ *δ* ≤-90°and (ii) 90°≤ *θ* ≤140° and 50°≤ *δ* ≤90°, which matched with the location of the PES scan minima (Figure 2b; Supplementary Figure S7a). The clusters represented the preferred direction of approach of the electrophile towards the lone-pairs of S. In the case of S···O contacts, a single cluster was observed at a different region of the *θ*-*δ* plot (115°≤ *θ* ≤155° and −50°≤ *δ* ≤50°), which overlapped with the PES scan minimum (Figure 2c; Supplementary Figure S7a). The direction corresponded to the approach of the nucleophile towards the σ-hole on S (Politzer *et al.*, 2013; Aakeroy *et al.*, 2019). Outliers in the plots were due to the presence of other strong interactions, such as other H-bonds and stacking interactions, within the molecules (Supplementary Figures S7b). In the CSD, the number of S···N interactions (**F4** in Figure 1b) was much less than S···O (**F3** in Figure 1b), possibly because of N being conjugated in most of the structures resulting in the lack of lone-pair electrons for the formation of Ch-bond.

### Delineation of Ch-bond from H-bond in groups having electrophilic and nucleophilic centers

We then proceeded to find the nature of bonding between S and functional groups having both electrophilic and nucleophilic centers, because such interactions are expected to be common in proteins. To address this, we studied the CSD for fragments **F5** and **F6** in which the electrophile and the nucleophile were separated by a single covalent bond (Figure 1a-b). A distance less than the sum of the van der Waals radii of S and O/N ensured that these fragments formed either S···H-O/N or S···O/N interaction, but, because of structural constraints, not both simultaneously. From this set of interactions, contacts satisfying *d_S···H_* ≤ 2.8 Å were assigned as H-bond and the rest as Ch-bond. Note that use of this filtering strategy excluded those H-bonds having the distance between S and O/N greater than the sum of their van der Waals radii, i.e. *d_S···H_* > 2.8 Å.

The CSD analysis revealed three clusters in the *θ*-*δ* plot (Figure 2d). Two of these clusters matched with those seen for the H-bond formed in **F1** and **F2** (Figure 2b). Most interactions in the two clusters satisfied the H-bond criterion of *d_S···H_* ≤ 2.8 Å (blue dots in Figure 2d; Supplementary Figure S7a). S···O/N interactions with *d_S···H_* > 2.8 Å (red dots in Figure 2d) primarily clustered at a region that matched with the cluster formed by **F3** and **F4** forming Ch-bond (Figure 2c). A few interactions in this cluster had *d_S···H_* ≤ 2.8 Å. Note that H-O/N groups that formed Ch-bond with S could also form H-bond with a neighboring acceptor atom (Supplementary Figure S7a).

In proteins, interactions of methionine or cystine with side chains of serine, threonine and tyrosine or the backbone amide or water (fragments **F7**-**F9** in Figure 1b) are equivalent to fragments **F5** and **F6**, and could result in either a Ch- or H-bond. As the coordinates of H in most crystal structures in the PDB are absent, we used the angular limits for *θ* and *δ* obtained by analyzing **F1**-**F4** to identify S-mediated H- and Ch-bonds in proteins. Due to this reason and because there were very few examples of X_1_-S-H in the CSD analysis discussed above, thiol group of cysteine was not included in this or any other analyses that follows.

Contacts were identified using the distance criteria of *d_S···O_* ≤ 3.32 Å and *d_S···N_* ≤ 3.35 Å. The *θ*-*δ* plot for **F7**-**F9** showed segregation of angular values into three clusters that corresponded to either H-bond (cyan boxes in Figure 2e and Supplementary Figure S6b, S8) or Ch-bond (red box in Figure 2e and Supplementary Figure S6b, S8). Thus, *θ* and *δ* allowed us to identify and distinguish Ch-bond from H-bond in protein structures. We note that the geometrical parameters for S-mediated H-bond and Ch-bond reported here (Figure 2a) can be used for modeling methionine and cystine in protein structures, particularly of those determined using low-resolution X-ray crystallography or electron cryo-microscopy data.

### Electronic environment of S determines the choice between Ch- and H-bond

We next sought to find what dictated the choice between formation of Ch-bond and H-bond. In general, the formation of H- or Ch-bond is observed to depend on the strength of lone-pairs and σ–holes on S, respectively (Adhikari and Scheiner, 2014; Kumar *et al.*, 2014). The magnitude of the lone-pairs and σ–holes are affected by the electronic properties of the substituent groups (Adhikari and Scheiner, 2014; Kumar *et al.*, 2014). Consequently, we studied the strength of lone-pairs and σ–holes of S using MESP in certain model systems that are relevant to biomolecules (Figure 3a). As expected, the MESP analysis revealed the presence of two V_min_ (MESP minimum) on S in all the model systems, which corresponded to the lone-pairs, and whose values were substituent dependent (Figure 3a). The electrostatic potential maps also showed the presence of two positive σ–holes along the extension of the S–X bonds except in the case of [Fe(SCH_3_)_4_] (Figure 3b). Strength of the σ–holes (V_s,max_) on S increased with the electron withdrawing power of the substituent and was negligible when it was chelated to metal. This confirmed that substituents on S modulated the electronic effects of the lone-pairs and σ–holes, which, consequently, was expected to affect the nature of the bond formed.

Next, we categorized all the structures in the CSD containing fragments listed in Figure 1b based on the substituents linked to S, i.e. S(Ar) = S in aromatic ring; M-S-M, M = any metal; M-S-Y, Y = any element except M; E-S-Y, E = any electron withdrawing group; R-S-R, R = saturated C, H or S. We analyzed these structures to check if S formed Ch-bond or H-bond (Figure 3c; Supplementary Table S2). The classification into Ch-bond or H-bond was based on the distance and angular criteria defined above. 87% of S(Ar) and 77% of E-S-Y formed Ch-bond (Figure 3c, Supplementary Table S2). In sharp contrast, more than 95% of M-S-M formed H-bond (Figure 3c, Supplementary Table S2). In comparison, saturated C/S/H substituents (R-S-R) appeared to have a lesser influence on the choice of the bond formed. The number of Ch-bond (62%) was slightly higher than H-bond (38%) (Figure 3c, Supplementary Table S2). The observations matched the expectations from the MESP analysis performed on representative molecules (Figure 3a-b). For example, the high occurrence of Ch-bond in S(Ar) and E-S-Y is correlated with their high V_max_ and low V_min_ values from the MESP calculations. Absence of a σ–hole in the MESP map of M-S-Y matched with the strong preference of metal-chelated S to form H-bond. Based on the above analysis, we concluded that if S is part of an aromatic ring or is bonded to an electron withdrawing group it is most likely to form a Ch-bond, while S coordinated with a metal can form H- but not Ch-bond.

### Disulfide-bonded S preferentially form Ch-bond while metal-chelated thiolate form H-bond

To find if protein structures in the PDB also display similar preferences, we studied interactions made by S of methionine or cystine with the hydroxyl, amino or carbonyl/carboxyl group of backbone amide, side chains of serine, threonine, tyrosine, aspartate, glutamate, arginine, lysine, histidine, asparagine, glutamine, tryptophan, and bound water (Supplementary Figure S9-S10 & Supplementary Table S3). Our analysis revealed that the disulfide-bonded Cys-S^γ^ was more frequently involved in Ch-bond (87%) than H-bond (13%). In comparison, Met-S^δ^ appeared to form H-bond (59%) only marginally more than Ch-bond (41%) (Figure 3d; Supplementary Table S4). The MESP analysis showed that S bonded to two methyl groups (C-S-C), as in methionine, had comparable values of V_min_ and V_s,max_ (Figure 3a-b), while V_s,max_ on a disulfide linked S (C-S-S) was larger than V_min_ (Figure 3a-b), thus providing a rationale for cystine to preferentially form Ch-bond.

Additionally, we also analyzed the interaction of aromatic S, which are often seen in ligands or drug molecules, with the above functional groups in proteins. Aromatic S preferentially formed Ch-bond with groups containing O (83%) consistent with MESP analysis (Figure 3d; Supplementary Table S4). This observation is also consistent with the previous reports that S in the aromatic rings of drugs containing thiophene, thiazole and thiadiazole groups interact with O in target proteins *via* Ch-bonds (Thomas *et al.*, 2015; Zhang *et al.*, 2015; Koebel *et al.*, 2016; Kristian *et al.*, 2018). S-mediated interaction with functional groups containing N did not show these features (Supplementary Table S4). This presumably is because the strongly delocalized lone-pairs of N in backbone amide or the side chain of arginine, lysine, histidine, asparagine, glutamine or tryptophan precluded formation of Ch-bond.

We next analyzed the PDB for non-covalent interactions formed by metal-chelated cysteines, which occur in many metalloproteins. Consistent with the rules stated above and independent of the identity of the metal, the thiolate of cysteine preferentially formed H-bond (Supplementary Table S4). As the resolution of the PDB structures analyzed were in general lower (resolution ≤ 2 Å) than the CSD structures, we relaxed the distance criterion for H-bond formation in proteins to *d_S···N_* ≤ 3.6 Å. In summary, our analyses of the structures in the CSD and the PDB revealed that the nature of interaction between functional groups containing electrophilic and nucleophilic centers and S was influenced by the substituent-dependent seesaw change in the strength of the lone-pairs and σ–holes on S (Figure 3e).

### Role of S in helix capping

Capping satisfies the H-bond forming abilities of the free backbone N-H or C=O of the terminal residues of an α-helix, and is important for the stability of α-helices in proteins and peptides (Aurora and Rose, 1998). The role of polar side chains of serine, threonine and asparagine, the acidic side chain of aspartate, the backbone amide of a neighboring residue, and metal-chelated S of cysteine in helix capping are well documented (Doig and Baldwin, 1995; Aurora and Rose, 1998), but not those of methionine and cystine. As the N-terminus and the C-terminus of α-helices have free backbone N-H (electrophile) and free backbone C=O (nucleophile), respectively, we asked if Met-S^δ^ or Cys-S^γ^ would interact and cap them. We analyzed protein structures in the PDB and found a number of examples of Met-S^δ^ or Cys-S^γ^ interacting with backbone amide at either the N-terminus or the C-terminus of α-helix (Figure 4a).

As reported previously (Doig and Baldwin, 1995), metal-chelated thiolates capped the N-terminus of the helix (Figure 4a,b; Supplementary Table S5). In contrast, 78% of Cys-S^γ^ capped the C-terminus by Ch-bond, while the remaining 22% capped the N-terminus (Figure 4b; Supplementary Table S5). Amongst the examples involving Met-S^δ^, 46% of the interactions were H-bonds with backbone N-H of the N-terminal residues and 54% Ch-bonds with backbone C=O of the C-terminal residues (Figure 4b; Supplementary Table S5).

### Augmentation of the stability of regular secondary structures by S

In addition to H-bond between *i* and *i*+4 residues and the capping interactions, other non-covalent interactions such as C-H···O and n→π^*^ are important for structural stability of α-helices (Manikandan and Ramakumar, 2004; Bartlett *et al.*, 2010b). Similarly, the structural stability of a β-sheet is dependent not only on inter-strand H-bonds between backbone O and N-H but also inter-strand C-H···O interactions (Derewenda *et al.*, 1995). An earlier study reported instances of methionine forming intra-helical and inter-strand Ch-bonds with backbone O (Pal and Chakrabarti, 2001). This prompted us to find if S could also contribute to the stability of the regular secondary structures, α-helices and β-strands, through H-bond and Ch-bond with backbone O or N-H.

We analyzed the PDB for interactions made by Met-S^δ^ or Cys-S^γ^ with backbone O or N-H of residues constituting α-helices and β-strands. In case of α-helices only internal residues were considered, because the interactions between residues at the helical termini and S were analyzed in the previous sections. Met-S^δ^ and Cys-S^γ^ formed Ch-bond with backbone O of α-helices and β-strands (Supplementary Figure S11, Supplementary Table S6). Examples of Ch-bonds with backbone N of α-helical and β-strand residues, though present, were considerably less in number (Supplementary Table S6). However, as would be expected, N-H···S bond was not observed in intra-helical regions, as the backbone N-H of *i^th^* residue was H-bonded to the backbone C=O of *i*−4^th^ residue.

Interestingly, we found N-H···S interaction in β-strand regions (Supplementary Figure S11 & Supplementary Table S6), many of which involved backbone N-H of β-strands at the edge of β-sheets (Figure 5a). The H-bond appeared to stabilize the free N-H. We also found Ch-bond formed by S with free backbone C=O of edge strands (Figure 5a). Many elements of negative-design that stabilize the edge strand of β-sheets have been documented previously, including interaction of other regions of the protein with the edge β-strand, disruption of backbone H-bond formation by proline or a β-bulge, or use of inward-pointing charged residues to prevent strand-mediated dimerization (Richardson and Richardson, 2002; Koga *et al.*, 2012). Our analysis revealed that H-bond or Ch-bond formed by backbone N-H or C=O of edge β-strand with a neighboring Met-S^δ^ or Cys-S^γ^ is another element of negative-design that can stabilize β-sheets (Figure 5a). Additionally, we found that in some proteins, the free backbone N-H of the insertion residue of a classical β-bulge formed H-bond with Met-S^δ^ located two residues ahead (Supplementary Figure S11).

To gauge the potential contribution of Ch-bond to lock protein conformation and stabilize regular secondary structures, we performed PES scan by varying the torsion angle *χ* about the covalent bond between C^α^ and C^β^ of the disulfide-bonded cysteine whose S formed a Ch-bond (Figure 5b-c). One fragment each from an α-helix and a β-strand were chosen for the calculations (Figure 5b). For ease of calculation, the side chain groups were excluded from the calculations (see Methods). A plot of relative conformational energy vs *χ* showed that the minimum conformational energy corresponded to the *χ* of the respective crystal structures (Figure 5c). The structures corresponding to the energy minimum were subjected to Atoms In Molecules (AIM) analysis which showed presence of Bond Critical Point (BCP) between S and O. *ρ* values of these BCPs were in the range suggested previously, i.e. 0.002-0.035 au (Bader, 1991). The analysis, thus, strongly suggested that Ch-bond could provide extra stability to a particular conformation in protein molecules.

### Ch-bond stabilizes β-turn containing motifs

Apart from stabilizing regular secondary structures, we found that S-mediated interactions stabilized β-turns in the motif CXXXXC in which cysteines at *i* and *i*+5 positions were linked by a disulfide bond. The motif, to be referred to as Case 1 (Figure 6a), adopted a unique conformation in which residues *i*+2 to *i*+5 formed a type II β-turn, with the backbone C=O of *i*+2 residue additionally forming a Ch-bond with the disulfide bonded S of the cysteine at *i*+5 position (Figure 6a; Supplementary Tables S7). The motif was found in diverse proteins, including members of the three-finger protein fold (for example PDB ID 6GBI), bacterial pore-forming toxin proaerolysin (PDB ID 1PRE), carboxylesterase Notum (PDB ID 4UZ1), lipoprotein lipase (PDB ID 6E7K), Type IV Pilins (PDB ID 5G24) and FAB (PDB ID 3QYC). Each of the six residues of the motif occupied a unique position in the Ramachandran plot (Figure 6b). The Ch-bond along with the H-bond between backbone C=O of *i*+2 and backbone N-H of *i*+5 residues, the latter being characteristic of a type II β-turn (Venkatachalam, 1968), appeared to stabilize the unique conformation of the motif.

The CXXXXC motif in the protein Cel7A had a type I β-turn instead of type II turn, which was immediately preceded by a type II’ β-turn (to be referred to as Case 2; Figure 6a,c; Supplementary Tables S7). The CXXXXC motif of ionotropic glutamate receptor (PDB ID 4YKI), α-conotoxin (PDB ID 1HJE), Xpd4 helicase (PDB ID 2VL7) and archeal prim-pol domain (PDB ID 1RO2) also had a conformation similar to Case 2 but without a preceding type II’ β-turn. A striking feature of the CXXXXC structural motifs in Cases 1 and 2 was the conservation of the Ramachandran angles of the constitutive residues (Figure 6b, c; Supplementary Table S7). In comparison, the Ramachandran angles of the six residues in the CXXXXC structural motifs lacking the Ch-bond were clearly less conserved. These structures could be divided into two groups - Case 3 had a type I β-turn and the corresponding H-bond between backbone C=O of *i*+2 and backbone N-H of *i*+5 residues (Figure 6a, d) and Case 4 which did not have any β-turn (Supplementary Table S7). This indicated that the Ch-bond played a role in the unique conformation adopted by the motifs.

To confirm this, we performed quantum chemical and AIM analyses of Cases 1 to 3 (for models refer Methods). Conformations of Cases 1 and 2 were found to be energetically favorable compared to the conformation in Case 3 (see E_*rel*_ values in Figure 6a). Interestingly, *ρ* values of BCP characterizing S···O bond and N-H···O bond were comparable in Case 1 (Figure 6e; Supplementary Table S8). This indicated similar strengths of H- and Ch-bonds, and demonstrated that the Ch-bond contributed significantly to the stability of the structural motif. In Case 2, in addition to the H-bonds, the Ch-bond also contributed to the stability of the structure (Figure 6e; Supplementary Table S8).

## Discussion

In this study, we have tried to understand the role of H- and Ch-bonds formed by divalent S in proteins. Computational analyses showed that the S-mediated interactions contributed to the stability of protein conformation and secondary structures. Hence, we conclude that S-mediated Ch- and H-bonds, like other weak interactions, are an important aspect of the energy landscape in protein folding that compensates for unfavorable conformational entropy change through favorable interactions (Grantcharova *et al.*, 2001; Dobson, 2003). Furthermore, we envisage that cooperativity among S-mediated interactions and other weak interactions is likely to modulate their strengths with direct implication to protein function, which remains to be studied. For example, we speculate that the propensity and strength of Ch-bond would increase upon delocalization of lone-pair electron density of Cys-S^γ^ to form an n→π^*^ interaction with a vicinal carbonyl group (Kilgore and Raines, 2018).

S-mediated Ch- and H-bonds can contribute to structural stability and substrate specificity of proteins, very much like other interactions formed by polar amino acids. However, S-mediated interactions can have properties different from other polar non-covalent interactions, for instance, the resistance of Ch-bond strength to solvent polarity (Pascoe *et al.*, 2017), thus bringing additional diversity to the repertoire of weak-interactions essential for biomolecular functions. This could be a reason why, despite their high biosynthetic cost (Doig, 2017), nature selected S-containing amino acids as part of the twenty building blocks of proteins. The wide variety of functionally relevant interactions made by S in proteins necessitates that these non-covalent interactions too are considered in the energy functions used for determining protein structures, folding pathways and binding properties. Also, the design and engineering of proteins and peptides would benefit from a better understanding of the distinct bonding properties of methionine and cysteine/cystine.

