## Supplementary Information for "Sulfur-mediated chalcogen versus hydrogen bonds in proteins: a seesaw effect in the conformational space"

Kayarat Saikrishnan[1*]

[1]Department of Biology, Indian Institute of Science Education and Research, Pune, 411008, India.

[2]Department of Chemical Sciences, Indian Institute of Science Education and Research, Mohali, 140306, India.

### Table of contents


| | content | Page |
|---|---|---|
| Figure S1 | Definition of the PES scan parameters | 03 |
| Figure S2 | A distribution of *d* values for all contacts in Figure 1b | 04 |
| Figure S3 | A distribution of *θ* values for all contacts in Figure 1b | 05 |
| Figure S4 | A distribution of *δ* values for all contacts in Figure 1b | 06 |
| Figure S5 | Definition of *ζ* | 07 |
| Figure S6 | *θ* vs *δ* plot for F2 & F4 with computationally calculated ΔEs | 08 |
| Figure S7 | Examples of contacts in F1-F6 along with outliers | 09 |
| Figure S8 | Examples of contacts in F7-F8 | 10 |
| Figure S9 | *θ* vs *δ* plots for all S⋯O contacts in PDB | 11 |
| Figure S10 | θ vs δ plots for all S⋯N contacts in PDB | 12 |
| Figure S11 | Examples of H-bond and Ch-bond interactions in α-helix and β-strand | 13 |
| Table S1 | Statistical details of all contacts mentioned in Figure 1b | 14 |
| Table S2 | Classification of CSD data based on electronic nature of S | 15 |
| Table S3 | Summary of the PDB data analysis | 16 |
| Table S4 | Classification of the PDB structures based on electronic nature of S | 18 |
| Table S5 | Summary of the PDB analysis investigating S in α-helix capping | 18 |
| Table S6 | Summary of the PDB analysis for Ch- & H- bond in α-helix and β-sheets | 19 |
| Table S7 | Summary of PDB analysis investigating CXXXXC motifs | 19 |
| Table S8 | Summary of AIM analysis for Cases 1-3 in Figure 6a | 20 |

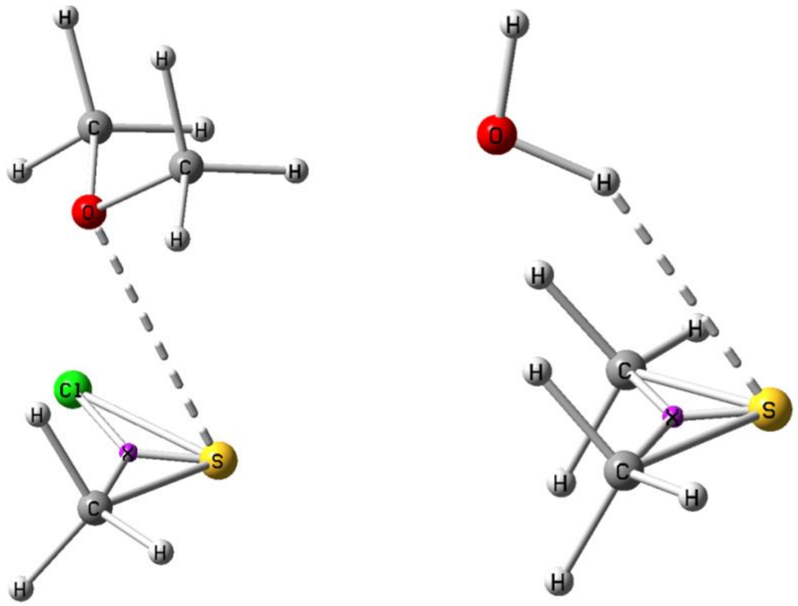


Figure S1. Definition of the PES scan parameters for some representative complexes corresponding to F1 and F3 are as follow; *d* (Distance between S and H/O/N) = constant *θ* (Angle between Z, S and H/O/N) = 60˚ to 178˚ (2˚ step) and *δ* (Dihedral angle between Z, S and H/O/N) = -90˚ to 90˚ (2˚ step). Similar definitions were used for F2 & F4. *d* values were obtained from energy minimized structures and kept constant during PES scan. *θ* and *δ* were varied in steps of 2˚. For $Cl(CH_3)S{:}N(CH_3)_3$ *θ* was varied from 90˚ to 178˚ instead 60˚ to 178˚ when -40˚ ≤ *δ* ≥ 40˚ to avoid steric clashes between atoms.

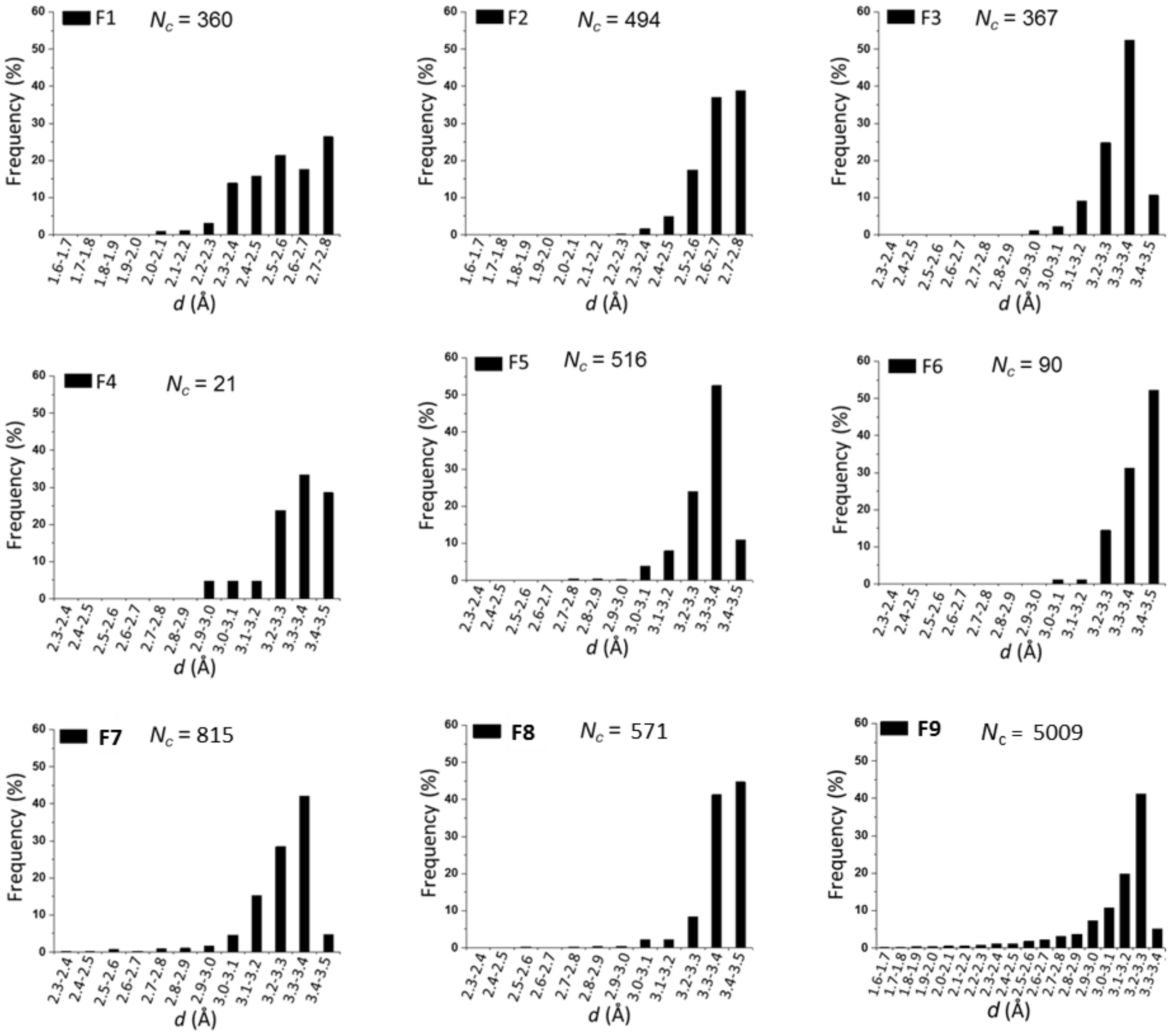


Figure S2. A distribution of *d* for all contacts mentioned in Figure 1b.

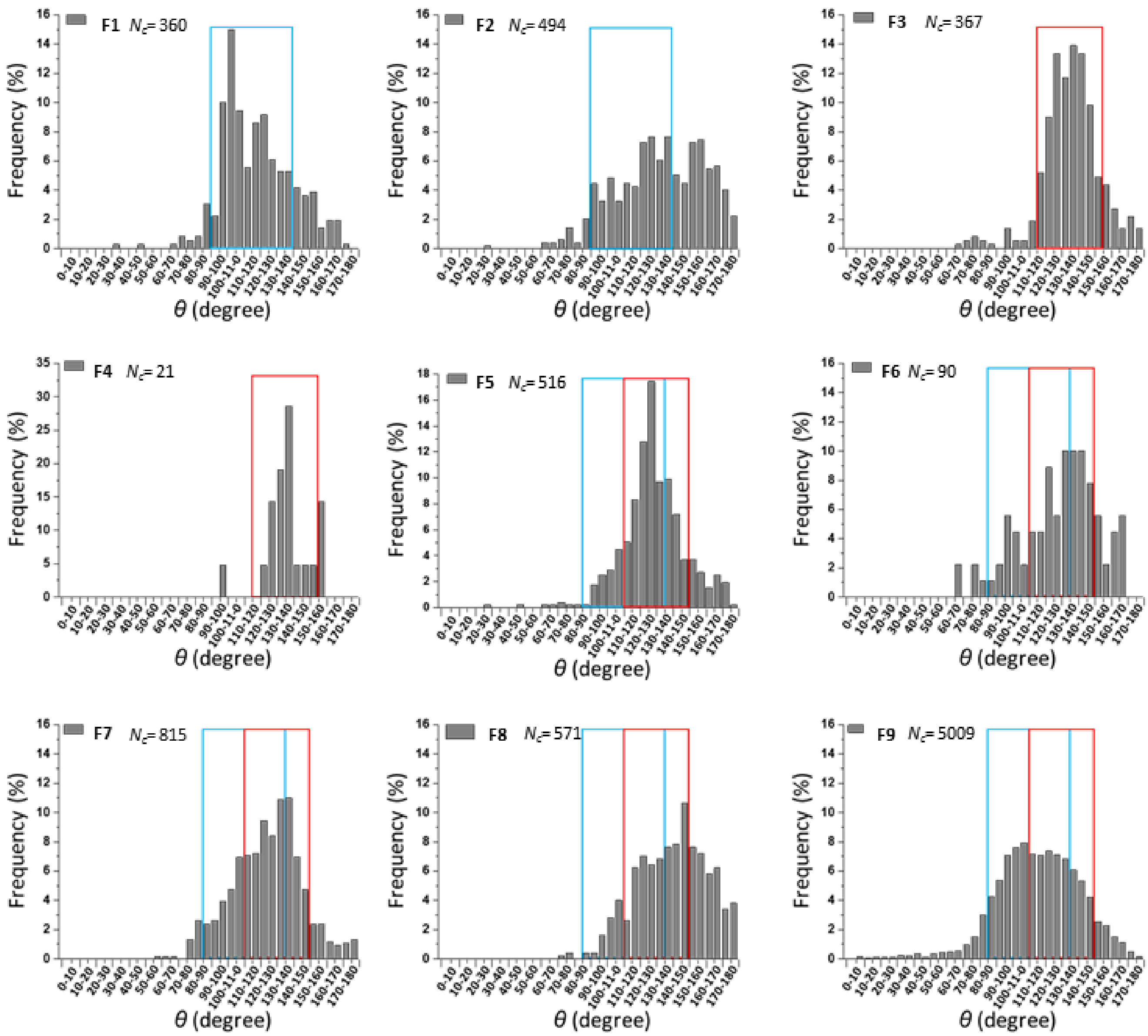


Figure S3. A distribution of $\theta$ values for all contacts mentioned in Figure 1b. Range of $\theta$ values used to investigate H-bond is marked in blue box while that for Ch-bond is marked in red box.

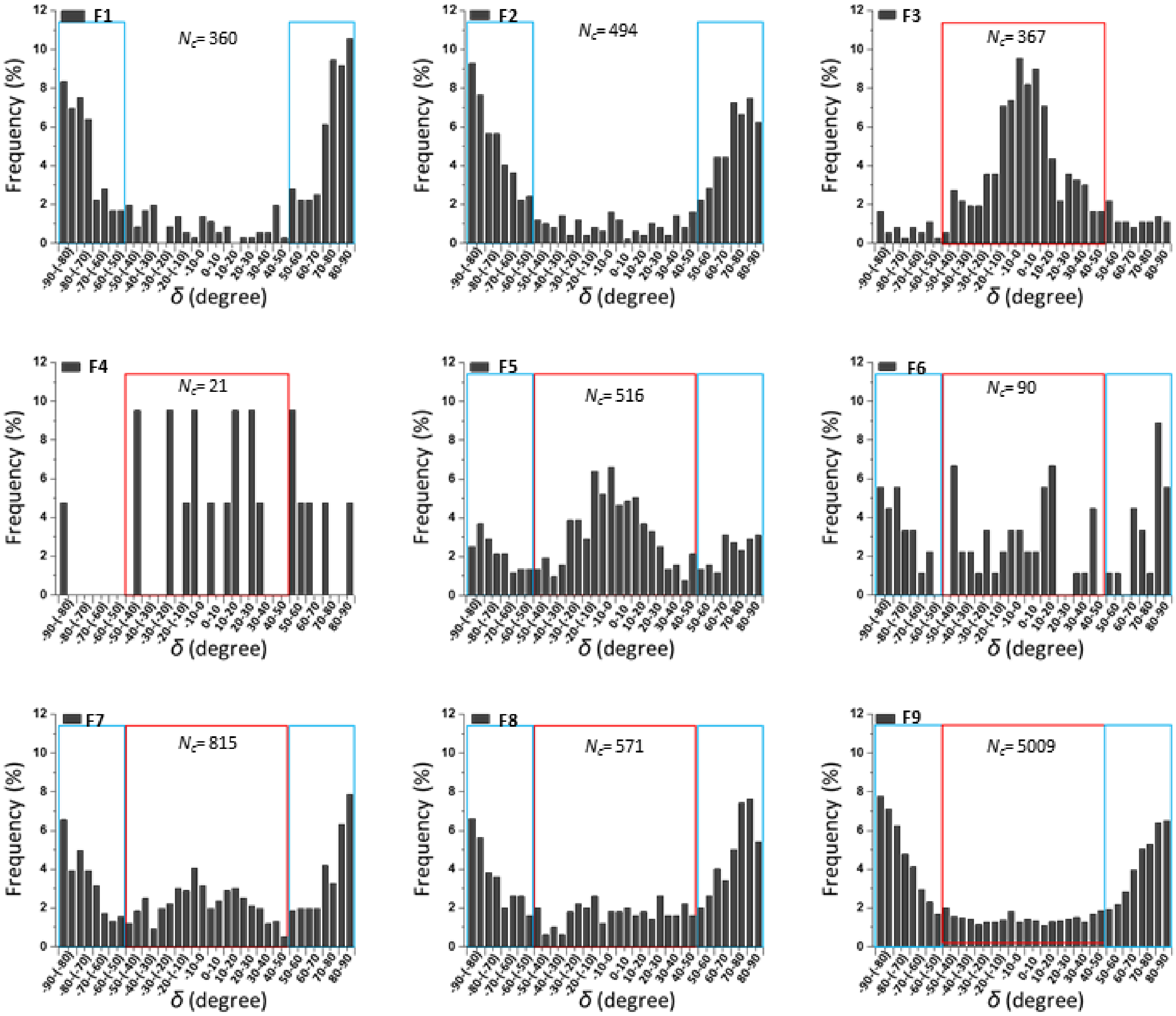


Figure S4. A distribution of $\delta$ values for all contacts mentioned in Figure 1b. Range of $\delta$ values corresponding to H-bond is marked in blue boxes while those for Ch-bond are in red box.

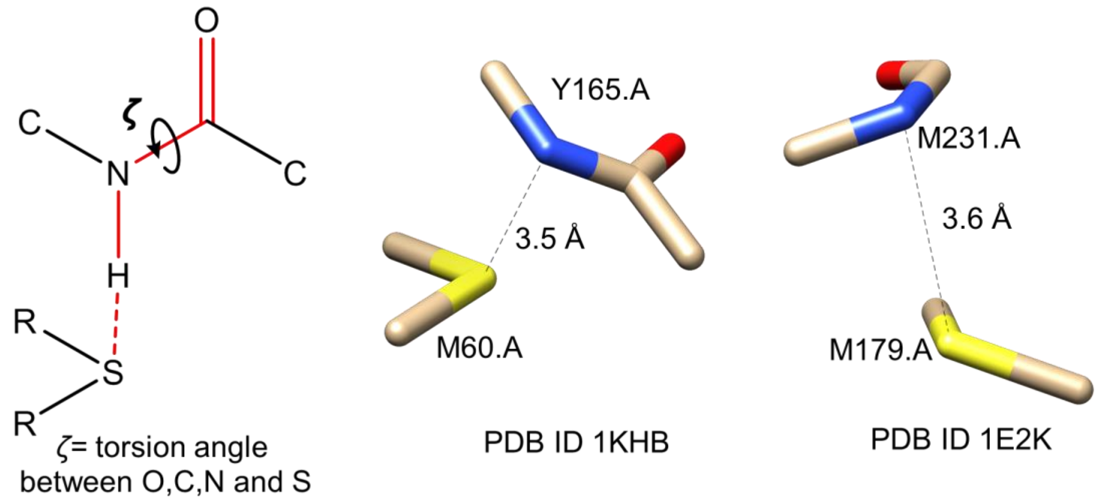


Figure S5. Definition of $\zeta$ along with some of the representative examples where $\zeta$ value was greater than 240° or less than 120°, which indicated that the respective H were pointing away from lone-pair region of S.

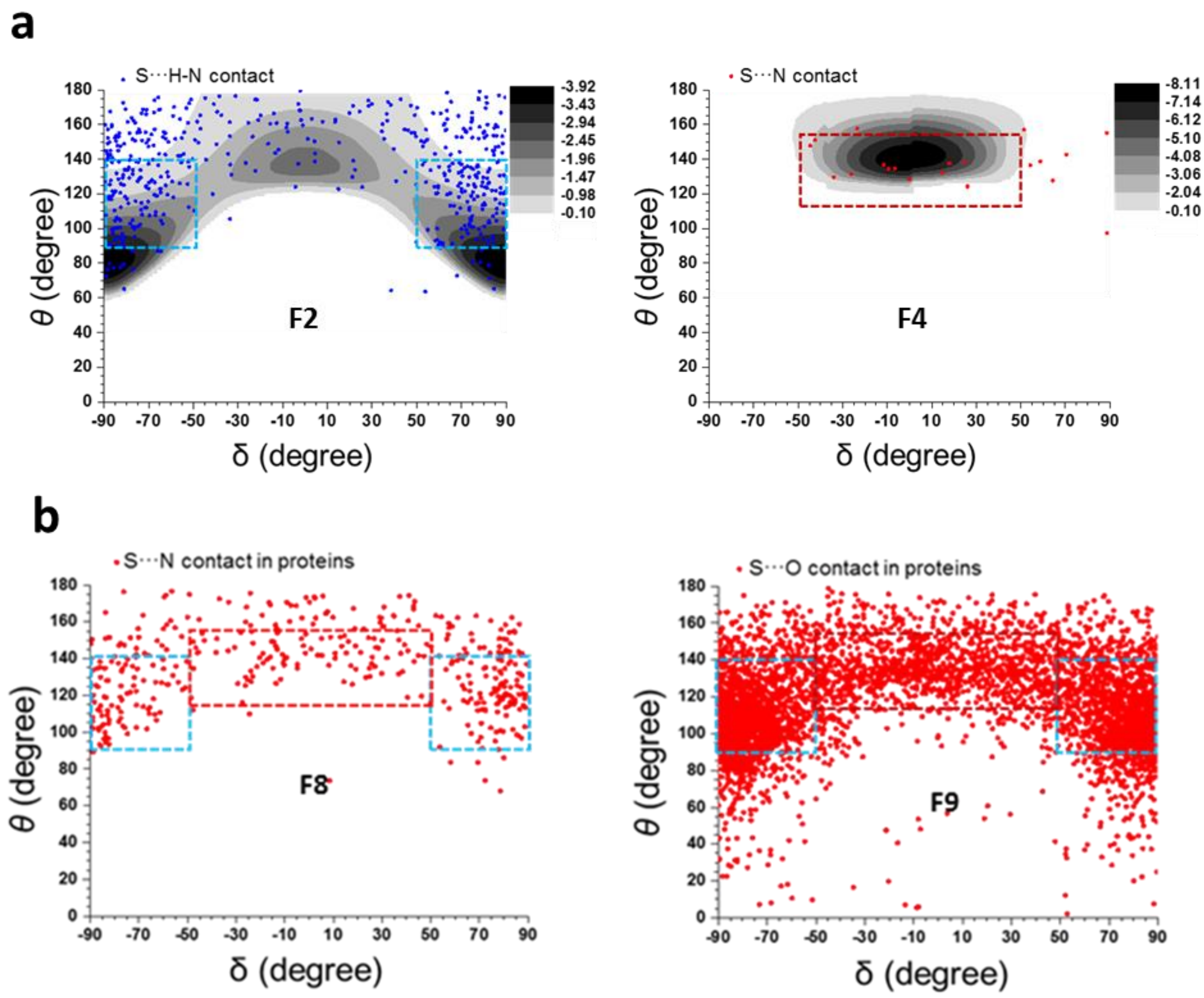


Figure S6. (**a**) Left panel: Mapping of $\theta$ and $\delta$ values of S···H-N contacts in fragment F2 with computationally calculated ΔEs in the background. $(CH_3)_2S{:}NH_3$ complex was used as a model system to calculate ΔEs. Right panel: For S···N contacts in fragment F4, $Cl(CH_3)S{:}N(CH_3)_3$ complex was used as model system to calculate ΔEs. (**b**) S···N (left panel) and S···O contacts (right panel) formed by methionine and cystine in fragments F8 and F9, respectively. In case of proteins, interacting atoms were separated by at least 6 covalent bonds.

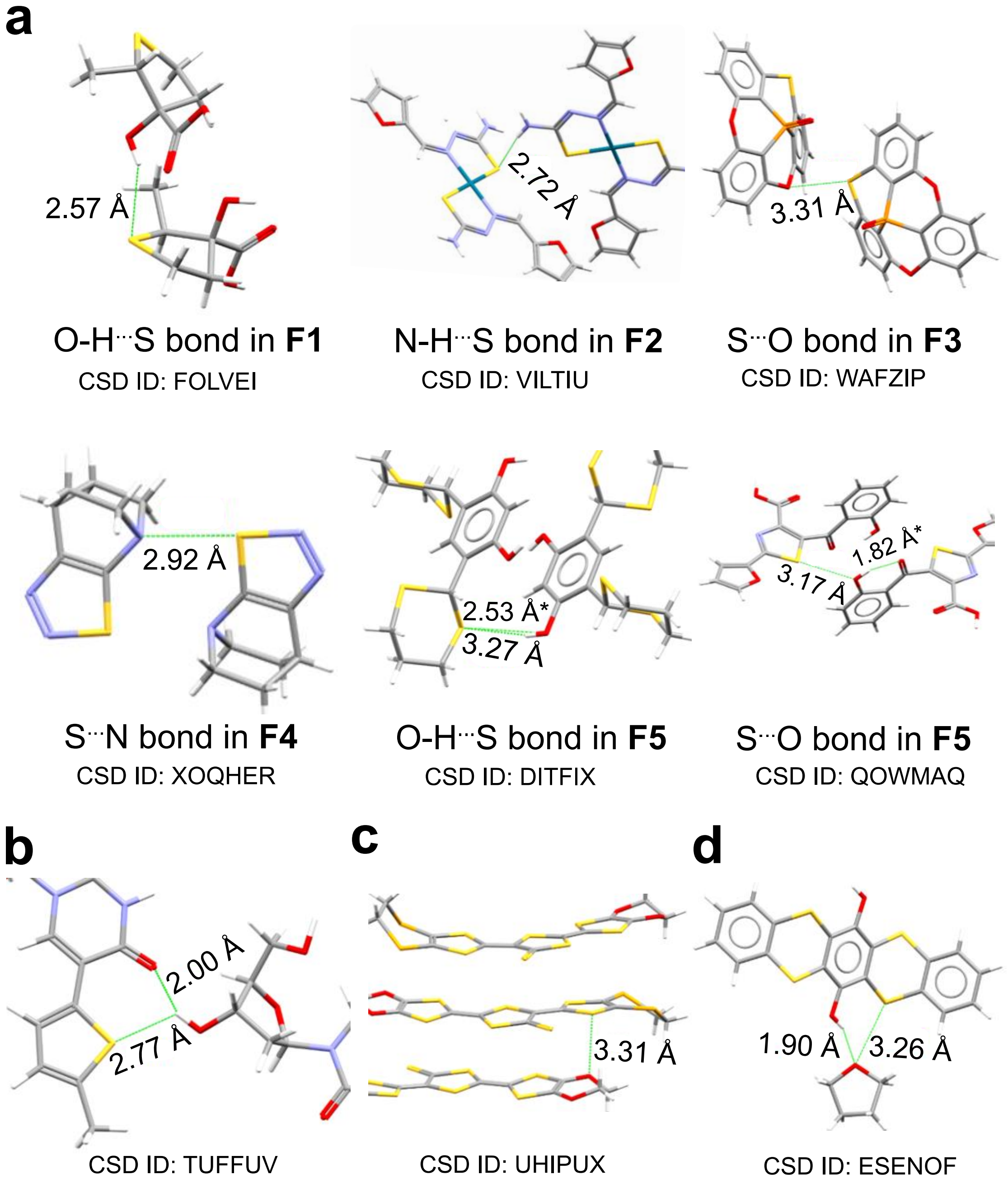


Figure S7. (**a**) Representative examples of H- and Ch-bond in F1-F5 with their CSD ID. In case of fragment F5, $d_{S\cdots H}$ is marked by an asterisk. Note that H-O/N groups that formed Ch-bond with S in F5 could form H-bond with a neighboring acceptor atom (**b**) Representative example of outliers for S···H-O contacts in F1 (**c**) Representative example of outliers of S···O contacts in F3 from those clustered around at $\theta$ =75˚ and $\delta$ =90˚ or -90˚ (refer Figure 1e) and (**d**) Representative examples of outliers of S···O contacts in fragment F3. All the examples are shown with their CSD ID

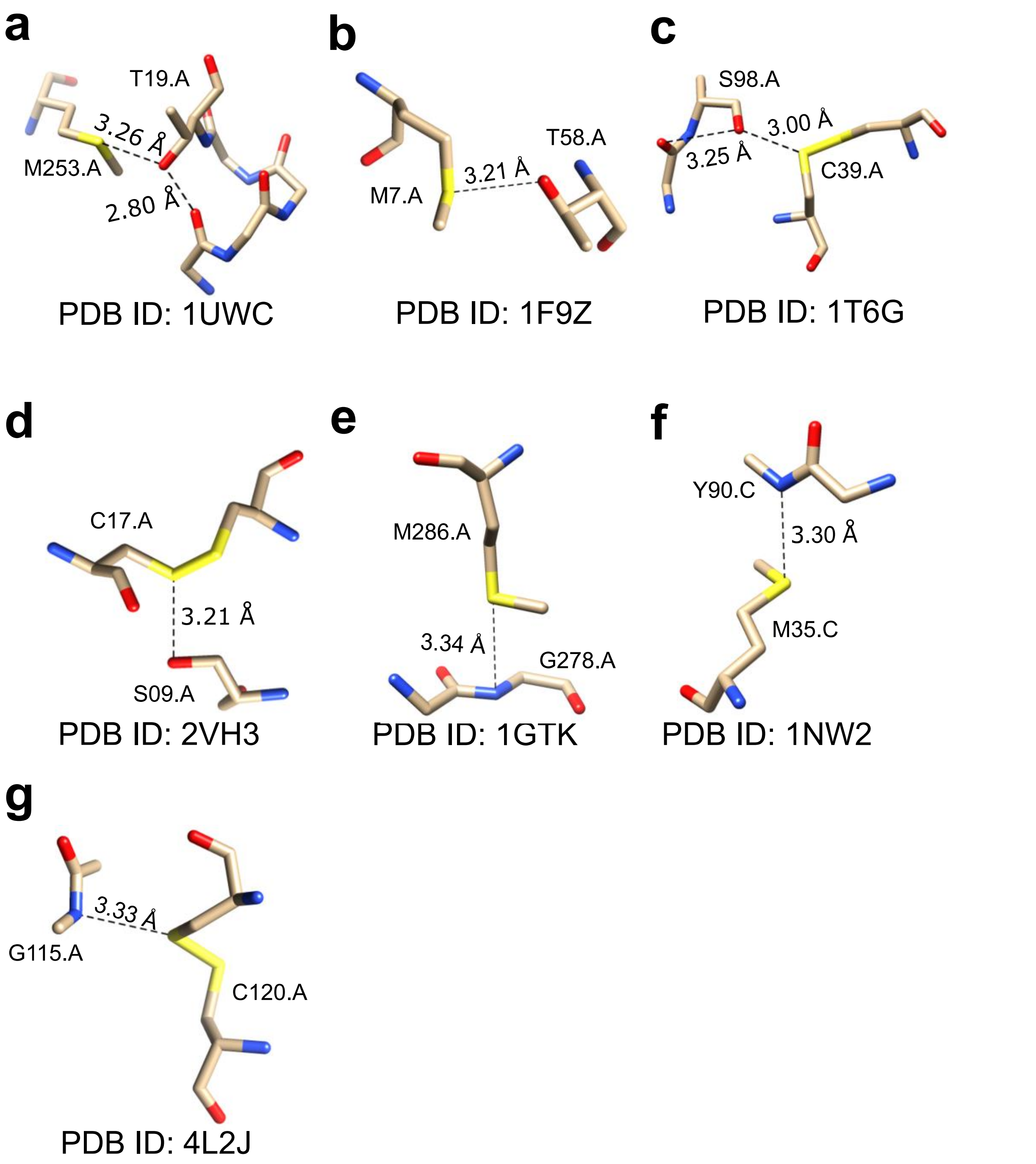


Figure S8. Representative examples of methionine-mediated **(a)** Ch-bond **(b)** H-bond in fragment F7. Representative examples of cysteine-mediated **(c)** Ch-bond **(d)** H-bond. Potential coexisting H-bonds are also illustrated. Representative examples of methionine-mediated **(e)** Ch-bond **(f)** H-bond in fragment F8 (**g**) Representative example of cysteine-mediated Ch-bond in F8. All structures are shown with their PDB IDs.

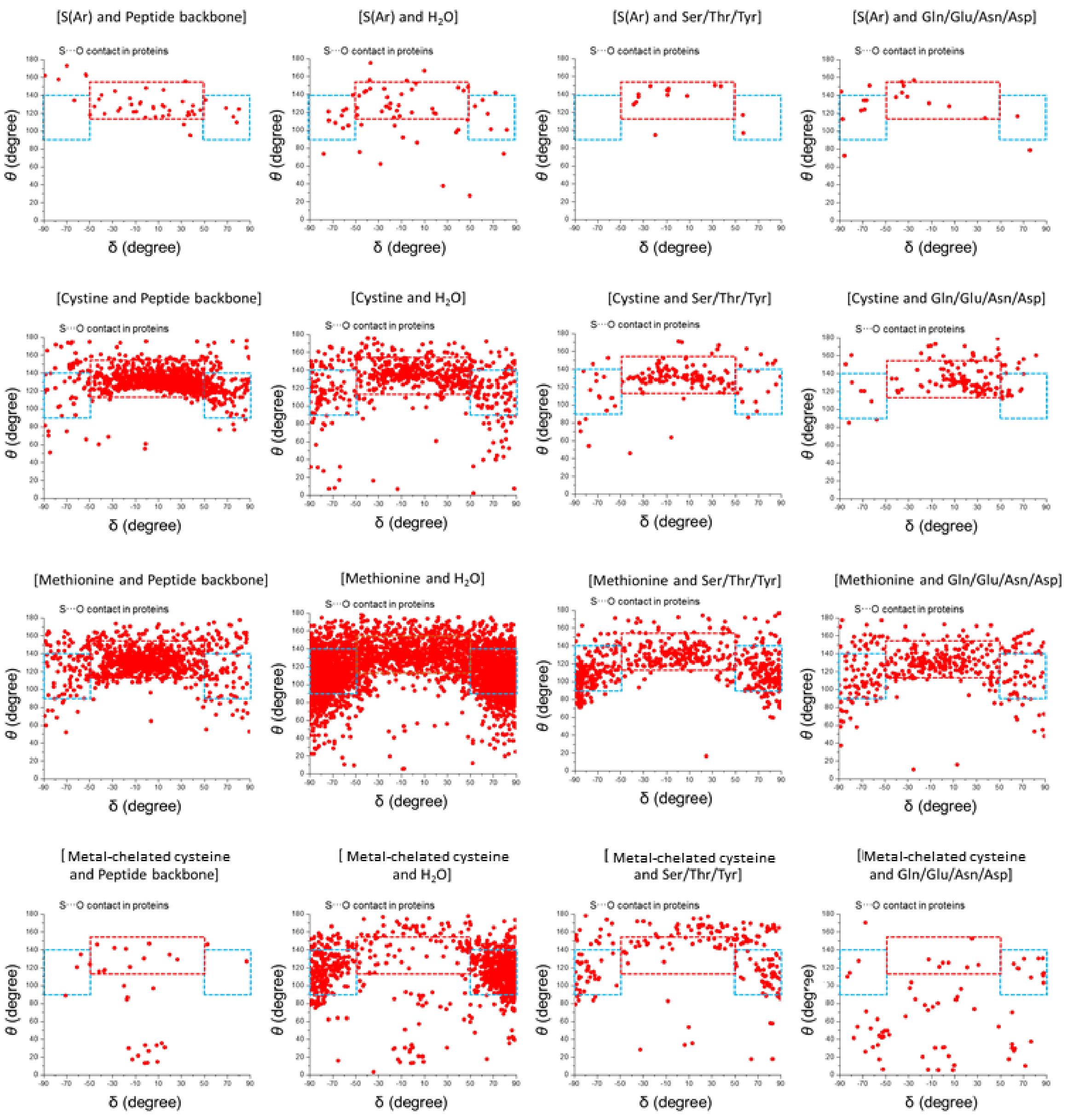


Figure S9: *θ*-*δ* plots for all S···O contacts in PDB (see Table S3). Resolution ≤ 2.0, Pair-wise sequence identity ≤ 90% and $R_{free}$ ≤ 25%.

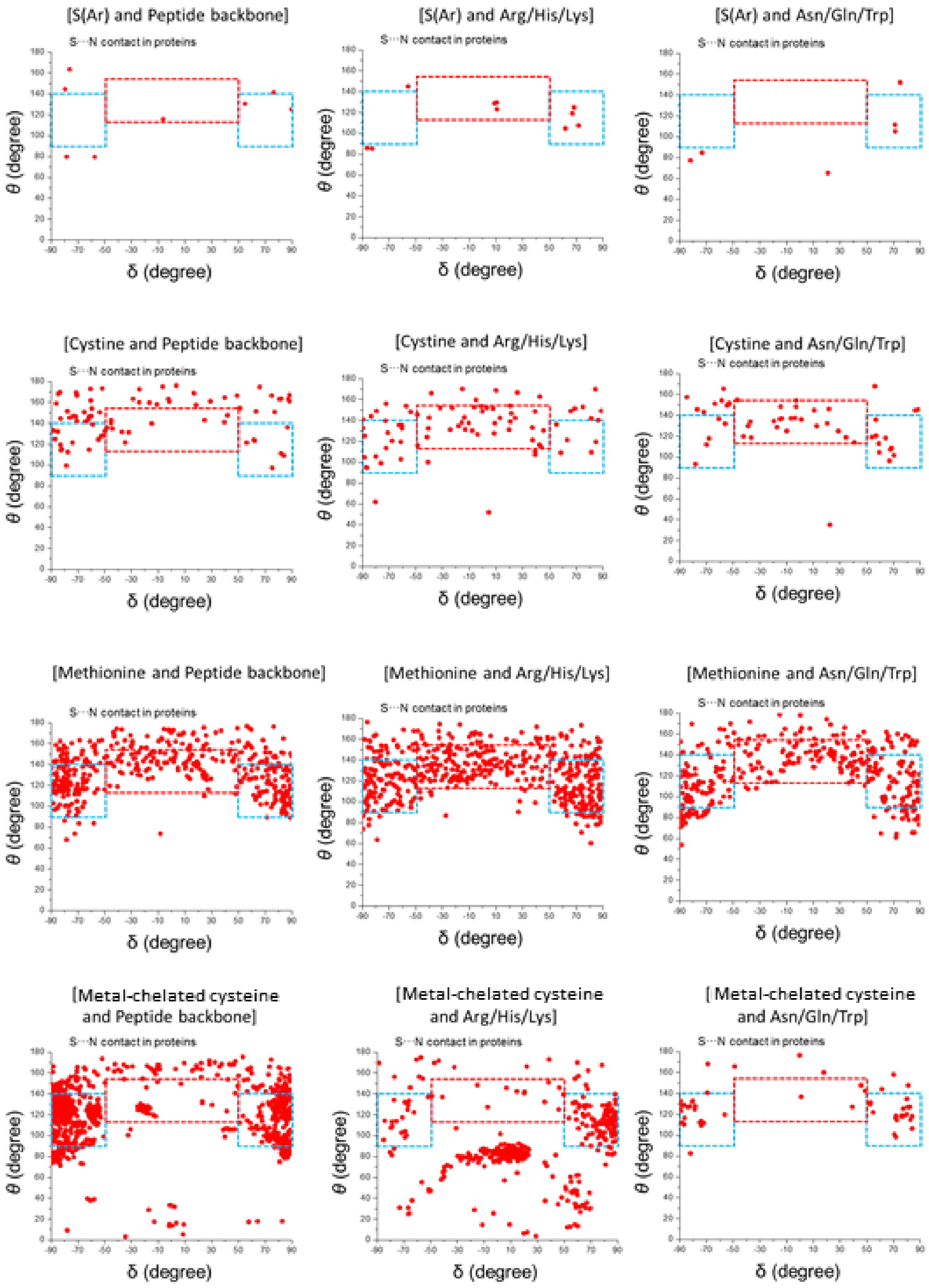


Figure S10: *θ*-*δ* plots for all S···N contacts in PDB (see TableS3). Resolution ≤ 2.0, Pair-wise sequence identity ≤ 90% and $R_{free}$ ≤ 25%.

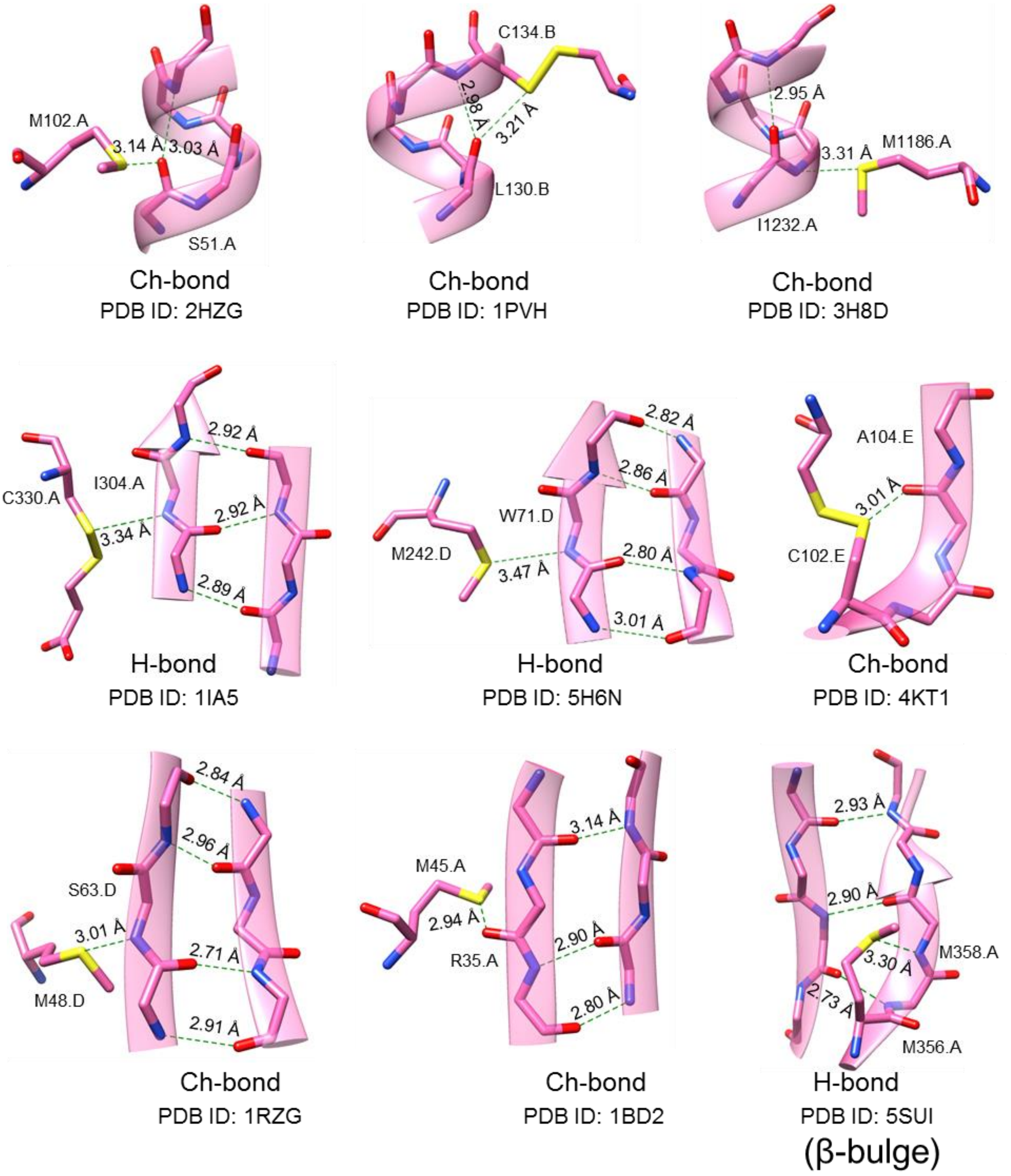


Figure S11. Representative examples of H-bonds and Ch-bonds formed by Met-S$^{\delta}$ and Cys-S$^{\gamma}$ with residues in α-helix and β-sheet. Also, an example of H-bond formed by Met-S$^{\delta}$ introducing β-bulge.

Table S1. Statistical details of all contacts mentioned in Figure 1b.

| Fragment | Criteria | Structural Parameter | Mean | Standard Deviation (± S.D.) |
|---|---|---|---|---|
| F1 | *d* ≤ 2.8 Å, 0° ≤ *θ* ≤ 180° -90° ≤ *δ* ≤ 90° | *d* | 2.56 Å | 0.16 Å |
| | | *θ* | 120.8° | 20.3° |
| | | *δ* | 4.0° | 71.3° |
| F2 | *d* ≤ 2.8 Å, 0° ≤ *θ* ≤ 180° -90° ≤ *δ* ≤ 90° | *d* | 2.66 Å | 0.10 Å |
| | | *θ* | 128.4° | 18.6° |
| | | *δ* | 0.2° | 69.6° |
| F3 | *d* ≤ 3.32 Å, 0° ≤ *θ* ≤ 180° -90° ≤ *δ* ≤ 90° | *d* | 3.21 Å | 0.09 Å |
| | | *θ* | 136.0° | 17.5° |
| | | *δ* | 3.6° | 34.3° |
| F4 | *d* ≤ 3.35 Å, 0° ≤ *θ* ≤ 180° -90° ≤ *δ* ≤ 90° | *d* | 3.22 Å | 0.1 Å |
| | | *θ* | 137.0° | 13.3° |
| | | *δ* | 13.0° | 44.1° |
| F5 | *d* ≤ 3.32 Å, 0° ≤ *θ* ≤ 180° -90° ≤ *δ* ≤ 90° | *d* | 3.21 Å | 0.10 Å |
| | | *θ* | 128.7° | 19.0° |
| | | *δ* | 0.1° | 47.8° |
| F6 | *d*1 ≤ 3.35 Å, 0° ≤ *θ* ≤ 180° -90° ≤ *δ* ≤ 90° | *d* | 3.28 Å | 0.08 Å |
| | | *θ* | 128.9° | 24.0° |
| | | *δ* | -1.4° | 58.7° |
| F7 | methionine, *d* ≤ 3.32 Å, 0° ≤ *θ* ≤ 180° -90° ≤ *δ* ≤ 90° | *d* | 3.19 Å | 0.13 Å |
| | | *θ* | 115.5° | 19.6° |
| | | *δ* | -6.0° | 65.5° |
| | cystine, *d* ≤ 3.32 Å, 0° ≤ *θ* ≤ 180° -90° ≤ *δ* ≤ 90° | *d* | 3.16 Å | 0.12 Å |
| | | *θ* | 128.5° | 18.6° |
| | | *δ* | -10.9° | 42,7° |
| F8 | methionine, *d* ≤ 3.35 Å, 0° ≤ *θ* ≤ 180° -90° ≤ *δ* ≤ 90° | *d* | 3.28 Å | 0.09 Å |
| | | *θ* | 134.5° | 21.4° |
| | | *δ* | 7.6° | 60.6° |
| | cystine, *d* ≤ 3.35 Å, 0° ≤ *θ* ≤ 180° -90° ≤ *δ* ≤ 90° | *d* | 3.27 Å | 0.12 Å |
| | | *θ* | 149.6° | 17.4° |
| | | *δ* | 14.6° | 56.9° |
| F9 | methionine, *d* ≤ 3.32 Å, 0° ≤ *θ* ≤ 180° -90° ≤ *δ* ≤ 90° | *d* | 3.09 Å | 0.28 Å |
| | | *θ* | 116.1° | 23.5° |
| | | *δ* | 3.3° | 68.3° |
| | cystine, *d* ≤ 3.32 Å, 0° ≤ *θ* ≤ 180° -90° ≤ *δ* ≤ 90° | *d* | 3.07 Å | 0.28 Å |
| | | *θ* | 93.7° | 52.1° |
| | | *δ* | -0.9° | 42.5° |

Table S2. Classification of the CSD data based on electronic nature of S. S-mediated H-bonds and Ch-bonds were identified using distance (*d*) and angular criteria (*θ* and *δ*) defined in the text.

| Fragment | S⋯H-O contacts ($N_c$) | S⋯H-N contacts ($N_c$) | S⋯O contacts ($N_c$) | S⋯N contacts ($N_c$) | Total Fragments ($N_f$) |
|---|---|---|---|---|---|
| M-S-M | 35 | 27 | 1 | 0 | 63 |
| M-S-Y | 172 | 136 | 15 | 3 | 326 |
| R-S-R | 32 | 15 | 68 | 9 | 124 |
| E-S-Y | 32 | 32 | 263 | 22 | 349 |
| S (Ar) | 11 | 18 | 236 | 22 | 287 |
| Total | 282 | 228 | 583 | 56 | 1149 |

Table S3: A summary of the number of interactions seen in PDB. H-bond and Ch-bond were identified using distance (*d*) and angular criteria (*θ* and *δ*) defined in the text.

| Interaction (Interacting residues) | $N_c$ | |
|---|---|---|
| | *Resolution ≤ 2.0 Å, pairwise sequence identity ≤ 90% and* $R_{free}$ *≤ 25%* | *Resolution ≤ 2.5 Å, pairwise sequence identity ≤ 90% and* $R_{free}$ *≤ 30%* |
| S···O contact [S(Ar) and Peptide backbone] | 51 | 92 |
| S···O contact [S(Ar) and Glu/Gln/Asn/Asp] | 20 | 31 |
| S···O contact [S(Ar) and Ser/Thr/Tyr] | 16 | 30 |
| S···O contact [S(Ar) and $H_2O$ ] | 63 | 114 |
| S···O contact [Cystine and Peptide backbone] | 1179 | 2041 |
| S···O contact [Cystine and Glu/Gln/Asn/Asp] | 140 | 265 |
| S···O contact [Cystine and Ser/Thr/Tyr] | 163 | 310 |
| S···O contact [Cystine and $H_2O$ ] | 671 | 1047 |
| S···O contact [Methionine and Peptide backbone] | 1059 | 2247 |
| S···O contact [Methionine and Glu/Gln/Asn/Asp] | 407 | 731 |
| S···O contact [Methionine and Ser/Thr/Tyr] | 652 | 1170 |
| S···O contact [Methionine and $H_2O$ ] | 4338 | 6221 |
| S···O contact [Metal chelated cysteine and Peptide backbone] | 38 | 67 |
| S···O contact [Metal chelated cysteine and Glu/Gln/Asn/Asp] | 82 | 138 |
| S···O contact [Metal chelated cysteine and Ser/Thr/Tyr] | 222 | 341 |
| S···O contact [Metal chelated cysteine and $H_2O$ ] | 1036 | 1377 |
| S···N contact [S(Ar) and Peptide backbone] | 8 | 18 |
| S···N contact [S(Ar) and Arg/His/Lys] | 10 | 18 |
| S···N contact [S(Ar) and Trp/Asn/Gln] | 6 | 9 |

| | | |
|---|---|---|
| S···N contact [cystine and Peptide backbone] | 95 | 229 |
| S···N contact [cystine and Arg/His/Lys] | 90 | 190 |
| S···N contact [cystine and Trp/Asn/Gln] | 55 | 107 |
| S···N contact [Methionine and Peptide backbone] | 476 | 965 |
| S···N contact [Methionine and Arg/His/Lys] | 565 | 1079 |
| S···N contact [Methionine and Trp/Asn/Gln] | 389 | 757 |
| S···N contact [Metal chelated cysteine and Peptide backbone] | 1153 | 2098 |
| S···N contact [Metal chelated cysteine and Arg/His/Lys] | 433 | 725 |
| S···N contact [Metal chelated cysteine and Trp/Asn/Gln] | 63 | 94 |

Table S4. Classification of the PDB data based on electronic nature of S. H-bonds and Ch-bonds were identified using the angular range of $\theta$ and $\delta$ defined in the text. Structures satisfying the criteria of resolution ≤ 2.0 Å, pairwise sequence identity ≤ 90% and $R_{free}$ ≤ 25% were used for the analysis.

| Fragment | S⋯H-O contacts ($N_c$) | S⋯H-N contacts ($N_c$) | S⋯O contacts ($N_c$) | S⋯N contacts ($N_c$) | Total Fragments ($N_f$) |
|---|---|---|---|---|---|
| M-S-C | 775 | 941 | 109 | 59 | 1884 |
| C-S-C | 2726 | 600 | 1958 | 390 | 5674 |
| C-S-S | 175 | 65 | 1462 | 77 | 1779 |
| S (Ar) | 17 | 8 | 82 | 4 | 111 |
| Total | 3690 | 1614 | 3611 | 530 | 9448 |

Table S5: A summary of the results of the PDB analysis performed to identify H-bond and Ch-bond formed by Cys-$S^{\gamma}$ or Met-$S^{\delta}$ that cap α-helices in proteins.

| Fragment | Total Number of α- helix capping contacts ($N_T$) [a] | | N-terminal α-helix capping contacts ($N_N$) [b] | | C-terminal α-helix capping contacts ($N_C$) [c] | |
|---|---|---|---|---|---|---|
| | *Resolution ≤ 2.0 Å* | *Resolution ≤ 2.5 Å* | *Resolution ≤ 2.0 Å* | *Resolution ≤ 2.5 Å* | *Resolution ≤ 2.0 Å* | *Resolution ≤ 2.5 Å* |
| C-S-S (Cystine) | 107 | 161 | 24 (22.4%) | 36 (22.4%) | 83 (77.6%) | 125 (77.6%) |
| C-S-C (Methionine) | 168 | 249 | 77 (45.8%) | 95 (38.2%) | 91 (54.2%) | 154 (61.8%) |
| C-S-M (Metal chelated cysteine) | 526 | 898 | 526 (100%) | 898 (100%) | 0 (0) | 0 (0) |

[a]Total number of S⋯O/ S⋯H-N contacts found capping α-helices. [b]Total number of S⋯H-N contacts found capping the N-termini of α-helices. [c]Number of S⋯O contacts found capping the C-termini of α-helices. The percentage in parenthesis was calculated using [($N_{N/C}$/ $N_T$) x100].

Table S6: A summary of the results of PDB analysis performed to identify H-bond and Ch-bond formed by Cys-$S^{\gamma}$ or Met-$S^{\delta}$ with residues in α-helix (only internal residues) and β-sheets. (Resolution ≤ 2.5 Å, pair-wise sequence identity ≤ 90% and $R_{free}$ ≤ 30%).

| Fragment | S···H-N interaction | | S···O interaction | | S···N interaction | |
|---|---|---|---|---|---|---|
| | α-helix | β- strand | α-helix | β- strand | α-helix | β- strand |
| C-S-S (Cystine) | 0 | 10 | 267 | 95 | 0 | 9 |
| C-S-C (Methionine) | 0 | 239 | 105 | 175 | 38 | 32 |

Table S7. A summary of the results of PDB analysis performed to study different CXXXXC motifs described in the text.

| Criteria | Number of contacts | |
|---|---|---|
| | For *Resolution* ≤ 2.5 Å, Pair-wise sequence identity ≤ 90% and $R_{free}$ ≤ 30% | For *Resolution* ≤ 2.0 Å, Pair-wise sequence identity ≤ 90% and $R_{free}$ ≤ 25% |
| Case 1<br>Ch-bond + H-bond<br>($d_{S\cdots O}$ ≤ 3.32 Å, 115˚≤$\theta$≤155˚, -50˚≤$\delta$≤50˚), ($d_{N5\cdots O2}$ ≤ 3.5 Å)<br>($d_{N4\cdots O1}$ > 3.5 Å)<br>Type II turn | 66 | 46 |
| Case 2<br>Ch-bond + H-bond<br>($d_{S\cdots O}$ ≤ 3.32 Å, 115˚≤$\theta$≤155˚, -50˚≤$\delta$≤50˚), ($d_{N\cdots O}$ ≤ 3.5 Å)<br>Type I and type II$^{I}$ turns | 07 | 05 |
| Case 3<br>Only H-bond<br>($d_{S\cdots O}$ > 3.32 Å, 115˚≤$\theta$≤155˚, -50˚>$\delta$>50˚ ($d_{N\cdots O}$ ≤ 3.5 Å)<br>Type I turn | 42 | 19 |
| Case 4<br>No Ch-bond or H-bond<br>($d_{S\cdots O}$ > 3.32 Å, 115˚>$\theta$>155˚, -50˚>$\delta$>50˚ ($d_{N5\cdots O2}$ > 3.5 Å)<br>($d_{N4\cdots O1}$ > 3.5 Å) | 171 | 97 |

Table S8. A summary of the results of AIM analysis for Case 1-3 in figure 5a,e.

| | Interaction No. | Interaction | ρ (kcal.mol-1) at BCP | E(SCF) In au | $E_{rel}$ (kcal.mol$^{-1}$) |
|---|---|---|---|---|---|
| Case1 (PDB ID 2FD6) | 1 | N-H···O | 0.02188 | -2009.27372 | -7.49 |
| | 2 | S···O | 0.02038 | | |
| | 3 | N-H···S | 0.01116 | | |
| | 4 | S···O | 0.00603 | | |
| Case2 (PDB ID 3CEL) | 1 | N-H···O | 0.01602 | -2009.27541 | -8.56 |
| | 2 | N-H···O | 0.01793 | | |
| | 3 | C-H···O | 0.01085 | | |
| | 4 | S···O | 0.00984 | | |
| Case3 (PDB ID 1HTR) | 1 | N-H···O | 0.01073 | -2009.26177 | 0.00 |
| | 2 | C-H···O | 0.00736 | | |
| | 3 | C-H···O | 0.00705 | | |
